# Human Stem Cell-Derived β-cells Expressing An Optimized CD155 Reduce Cytotoxic Immune Cell Function for Application in Type 1 Diabetes

**DOI:** 10.1101/2025.08.12.669867

**Authors:** Matthew E. Brown, Jessie M. Barra, Marcus R. Pina, James Proia, Todd M. Brusko, Holger A. Russ

**Affiliations:** Diabetes Institute, University of Florida, Gainesville, FL, USA; Department of Pathology, Immunology and Laboratory Medicine, College of Medicine, University of Florida, Gainesville, FL, USA; Department of Pharmacology and Therapeutics, College of Medicine, University of Florida, Gainesville, FL, USA; Department of Pediatrics, College of Medicine, University of Florida, Gainesville, FL, USA; Department of Biochemistry and Molecular Biology, College of Medicine, University of Florida, Gainesville, FL, USA

**Keywords:** Type 1 diabetes, autoimmunity, stem cell-derived β-like cell, TIGIT, CD155

## Abstract

Insulin-producing β-cell replacement therapies offers a potential treatment for type 1 diabetes (T1D) but faces challenges from donor shortages and immune rejection. Stem cell-derived β-cells (sBC) provide a renewable source but remain vulnerable to immune attack. We engineered human pluripotent stem cells to express either the wildtype (WT) or a high-affinity mutant (Mut) variant (rs1058402, G>A; Ala67Thr) of the NK and T cell checkpoint inhibitor CD155 before differentiation into sBC. Modified sBC maintained upregulated CD155 expression and showed enhanced binding to co-receptor ligands. Co-culture studies revealed that CD155 Mut-expressing sBC suppressed CD8^+^ T cell and NK cell activation and proliferation by preferentially engaging the co-inhibitory receptor TIGIT. Both CD155 Mut sBC lines reduced autoreactive CD8^+^ T cell- and NK cell-mediated sBC destruction and cytotoxic molecule secretion. This protection was lost with TIGIT blockade, confirming the role of CD155-TIGIT signaling in antagonizing immune cell-mediated killing. Our findings suggest that high-affinity CD155 expression enhances immune evasion of sBC, improving their potential for restorative therapy in T1D.

**Teaser:** Engineered β-cells with a mutant CD155 help evade immune attack, offering a promising therapeutic approach for type 1 diabetes.

## Introduction

Pancreatic β-cell replacement strategies hold great promise as a treatment option for patients with type 1 diabetes (T1D). However, challenges for the widespread implementation of this potentially curative approach include limited cadaveric donor tissue availability, immediate side-effects and long-term complications of systemic immune suppression, and cellular stress- or immune-mediated graft failure in a majority of patients by five years post-infusion (1–5). The differentiation of human stem cell-derived β-cells (sBC) from human pluripotent stem cells (hPSC) negates the requirement for organ donation, yielding a potentially limitless pool of human β-cells for clinical use (6, 7). Improvements in differentiation protocols over the past decade now allow for the generation of functional, glucose-responsive sBC that can restore glycemic control in pre-clinical animal models (8–14) and in an ongoing first Phase I/II clinical trial in human patients with T1D (clinicaltrials.gov; NCT04786262).

Still, similar to cadaveric islets, sBC are susceptible to immune-mediated destruction upon transplantation, necessitating the use of systemic immunosuppressive drugs, thereby excluding a larger patient population, due to the potential risk of serious secondary complications, such as viral infections, nephrotoxicity, and neurotoxicity (15, 16). To enhance the widespread implementation of sBC therapies, alternative approaches to prevent immune-mediated graft destruction are essential. Patients with T1D can mount two unique immune cell responses toward transplanted β-cells: 1) allogeneic rejection of the immunologically mismatched tissues, and 2) a recurrence of autoimmunity toward β-cell autoantigens. Both immune responses must be efficiently suppressed to maintain the long-term survival and function of any sBC graft.

With advances in genome engineering, hPSC can now be effectively modified to modulate the expression of key immune regulatory genes prior to differentiation into sBC (17–22). Multiple groups have demonstrated that this approach is efficacious in suppressing allogeneic and potentially even xenogeneic immune responses toward transplanted sBC in pre-clinical models (17, 20, 23). However, there is limited understanding of whether any modification can withstand autoimmune attacks, largely due to the lack of appropriate human model systems.

The immune checkpoint ligand and adhesion molecule, CD155 (poliovirus receptor; PVR), modulates immune responses through interactions with the NK cell and T cell co-inhibitory receptor, TIGIT (T cell immunoreceptor with Ig and ITIM-like domains) (24). Although CD155 can also participate with CD226 (DNAM-1; K_d_=119 nM) and CD96 (TACTILE; K_d_= 37.6 nM) to promote co-stimulatory and co-inhibitory signaling, respectively, CD155 preferentially interacts with TIGIT (K_d_ = 3.15 nM) to restrain T cell activation (25).

In the context of autoimmunity, there has been increased interest in leveraging this signaling axis to restore immune tolerance by skewing NK cells and T cells away from CD155/CD226 signaling and toward CD155/TIGIT signaling, given the presence of a T1D-risk associated single nucleotide polymorphism (SNP) in *CD226* (rs763361; C>T), thought to enhance downstream signaling (26). We demonstrated that a conditional knockout (cKO) of *Cd226* in regulatory T cells (Tregs) reduces disease incidence in the non-obese diabetic (NOD) model of autoimmune diabetes by increasing the expression of TIGIT on pancreatic CD4^+^ T cells (27) and, most recently, that monoclonal antibody blockade (mAb) of CD226 reduces disease incidence in the NOD by enhancing the immunoregulatory function of Tregs and inhibiting the function of pro-inflammatory effector T cells (28). Importantly, we have recently reported that pancreatic β-cells exhibit reduced expression of CD155 during T1D pathogenesis, suggesting dysregulation within this signaling pathway (29). Further, approaches seeking to bolster CD155/TIGIT signaling to limit T cell activity have shown promise in the treatment of autoimmunity, including the use of recombinant TIGIT-Immunoglobulin (TIGIT-Ig) fusion proteins to prolong survival in the NZB/W F1 mouse model of lupus (30), as well as the expression of TIGIT on CD4^+^ T cells using lentivirus to reduce collagen-induced rheumatoid arthritis (RA) in a BALB/C mouse model (31).

In oncology, the CD155/TIGIT checkpoint has been characterized as a critical mechanism for tumor immune escape by disrupting co-stimulatory signaling (32) and inducing a loss of effector function in tumor-infiltrating T cells. Consistent with these findings, the expression of a high-affinity mutant (rs1058402; G>A; Ala67Thr) of CD155 in patients with small-cell lung cancer demonstrated poorer treatment outcomes (33). Biolayer interferometry experiments revealed this mutant CD155 exhibits greater binding affinity towards both TIGIT and CD226 (34). Furthermore, natural killer (NK) cells co-cultured with a mutant CD155 (CD155 Mut)-expressing epithelial line exhibited reduced activation, compared to NK cells co-cultured with the wild-type CD155 (CD155 WT)-expressing epithelial line (34).

To understand whether the expression of either the CD155 Mut or CD155 WT could reduce allogenic and antigen-specific, autoreactive T cell responses as well as NK cell responses, we differentiated genome-edited hPSC lines into functional sBC and performed detailed mechanistic *in vitro* co-culture studies with islet antigen-reactive primary human T cells and NK cells. We investigated the immunogenicity of each human sBC line and how the modulation of the CD155 signaling pathway may confer protection from allo- and autoimmunity to better inform future pancreatic β-cell replacement therapies.

## Materials & Methods

### Generation of WT and Mutant CD155 Constructs

CD155 plasmid template was purchased from OriGene (RC202254) followed by transformation, amplification, and purification in NEB 5-alpha Competent *E. Coli.* To generate the Ala67Thr mutant version of CD155, Q5 site-directed mutagenesis (NE Biolabs, E0554S) was performed followed by transformation, amplification, and purification as with the original template. Confirmation of the mutation was performed through Sanger Sequencing. Both WT and Mut CD155 constructs were then amplified through PCR and ligated into a targeting backbone (made in-house) with homology arms for the endogenous AAVS1 locus, neomycin resistance gene, and a CAG promoter prior to nucleofection into hPSC (described below). All primers used are listed in **Table S1**.

### hPSC Culture and TALEN-mediated Engineering of CD155 WT and CD155 Mut Lines

The undifferentiated hPSC Mel1^INS-GFP^ line (NIH registry #0139) (35) was dissociated into single cells using TrypLE incubation at 37°C for 6 min. Digestion was then quenched with mTeSR+ media and live cells counted using a Countess 3 cell counter (ThermoFisher Scientific). 2×10^6^ cells were transferred into microcentrifuge tubes and washed with PBS. Washed cells were then prepared for nucleofection of TALEN mediated knock-in (KI) of either a WT CD155 or Ala67Thr point mutated (Mut) CD155 gene under control of a CAG promoter into the endogenous AAVS1 locus (**Supplemental Figure 1A, 1B**). Cells were nucleofected in P3 buffer per the Amaxa P3 Primary cell 4D-Nucleofector kit protocol (V4XP-3024) using the CB-150 program. AAVS1-TALEN-L, AAVS1-TALEN-R (Addgene plasmid #59025 and 59026), and CAG-CD155 expression plasmids (generated in-house) were nucleofected into hPSC Mel1^INS-GFP^ cells.

Nucleofected cells were then plated in 10cm plates with 10 μM ROCK inhibitor (Y-27632, R&D Systems #1254-50) and SCR7 (Excess Bioscience #M60082). After 24 hours, neomycin selection (50 μg/ml) was performed for 6 days. Remaining colonies were picked and expanded for further characterization. Genomic DNA was extracted from targeted colonies and PCR analysis (**Supplemental Figure 1C** and **Table S1**) and flow cytometry for TALEN KI was performed to identify successfully modified clones. Once edited clones of CD155 WT and CD155 Mut expressing hPSC were identified, the predominant clone utilized for the majority of further studies as well as the parental Mel1 hPSC line were sent for karyotyping via G-banding analysis (UF Health Pathology Cytogenetics Core). All submitted samples came back karyotypically normal (**Supplemental Figure 1D-F**).

### sBC Differentiation

hPSC Mel1^INS-GFP^, CD155 WT, or CD155 Mut stem cell lines were maintained on human embryonic stem cell (hESC) qualified Cultrex (Biotechne #3434-005-002) in mTeSR+ media (STEMCELL Technologies #05826). Differentiation to stem cell-derived β-like cells (sBC) was carried out in a suspension-based, magnetic stirring system (Reprocell #ABBWVS03A-6, #ABBWVDW-1013, #ABBWBP03N0S-6), as previously described (9, 19, 20). Briefly, 90% confluent hPSC cultures were dissociated into single-cell suspensions by incubation with TrypLE for 6 min (Gibco #12-604-021). Live cells were counted using a Countess 3 cell counter (ThermoFisher Scientific), followed by seeding 0.5 × 10^6^ cells/mL in mTeSR+ media supplemented with 10 μM ROCK inhibitor in bioreactors. 3D cluster formation was performed for 48 hours followed by washing of clusters with RPMI twice and induction of definitive endoderm (DE) differentiation using d1 media [RPMI containing 0.2% FBS, 1:5,000 ITS (Gibco #41400-045), 200 ng/ml Activin A (R&D Systems #338-AC-01M), and 3 μM CHIR99021 (STEMCELL Technologies #72054)]. Differentiation media was changed daily by letting clusters settle by gravity for 3-10 min. Most supernatant was removed by aspiration; fresh media was added, and stirrer flasks were placed back on the system. sBC differentiation was based on our published protocol (36) with modifications as outlined below. Differentiation medias are as: **day 2-3**: RPMI containing 0.2% FBS, 1:2,000 ITS, and 100 ng/ml Activin A; **d4-5**: RPMI containing 2% FBS, 1:1,000 ITS, and 50 ng/ml KGF (Peprotech #100-19-1MG); **d6**: DMEM with 4.5 g/L D-glucose (Gibco #11960-044) containing 1:50 N-21 MAX (Biotechne #AR008), 1:100 NEAA (Gibco #11140-050), 1mM Sodium Pyruvate (Gibco #11360-070), 1:100 GlutaMAX (Gibco #35050-061), 3 nM TTNPB, (R&D Systems #0761), 250 nM Sant-1 (R&D Systems #1974), 250 nM LDN (STEMCELL Technologies #72149), 30 nM PMA (Sigma Aldrich #P1585-1MG), 50 μg/ml 2-phospho-L-ascorbic acid trisodium salt (VitC) (Sigma #49752-10G); **d7**: DMEM containing 1:50 N-21 MAX, 1:100 NEAA, 1 mM Sodium Pyruvate, 1:100 GlutaMAX, 3 nM TTNPB, and 50 μg/ml VitC; **d8-9**: DMEM containing 1:50 N-21 MAX, 1:100 NEAA, 1 mM Sodium Pyruvate, 1:100 GlutaMAX, 100 ng/ml EGF (R&D Systems #236-EG-01M), 50 ng/ml KGF, and 50 μg/ml VitC; **d10-15**: DMEM containing 1:50 N-21 MAX, 1:100 NEAA, 1 mM Sodium Pyruvate, 1:100 GlutaMAX, 10 μg/ml Heparin (Sigma #H3149-250KU), 2 mM N-Acetyl-L-cysteine (Cysteine) (Sigma #A9165-25G), 10 μM Zinc sulfate heptahydrate (Zinc) (Sigma #Z0251-100g), 1x BME, 10 μM Alk5i II RepSox (R&D Systems #3742/50), 1 μM 3,3’,5-Triiodo-L-thyronine sodium salt (T3) (Sigma #T6397), 0.5 μM LDN, 1uM Gamma Secretase Inhibitor XX (XXi) (AsisChem #ASIS-0149) and 1:250 1 M NaOH to adjust pH to ∼7.4; **d16-30**: CMRL (Gibco #11530-037) containing 1:50 N-21 MAX, 1:100 NEAA, 1:100 GlutaMAX, 10ug/ml Heparin, 2mM Cysteine, 10 μM Zinc, 1x BME, 1 μM T3, 10 μM Alk5i II RepSox, 50ug/ml VitC, 1:1000 Trace Elements A (Corning # 25-021-CI),1:1000 Trace Elements B (Corning # 25-022-CI) and 1:250 NaOH to adjust pH to ∼7.4. All media also contained 1x Penicillin-Streptomycin.

### Cryopreservation and Thawing of sBC

Day 23 sBC were dissociated into single cells and cryopreserved as described (9, 19). Briefly, clusters were digested and filtered using a cell strainer into FACS 5 mL tubes. Cells were counted and resuspended at 3 x 10^6^ cells/100 μL of CryoStor® CS10 (StemCell Technologies). Cells were placed in cryovials (100 μL/vial), transferred into pre-cooled freezer buddies, and placed into the -80°C overnight before transfer to liquid nitrogen for long-term storage. For thawing, frozen vials of sBC were placed in a 37°C bead bath for 3 minutes to warm. Then, 1 mL of warm sBC media (d16-30 media described above) was added to the cryovial dropwise before the entire volume was transferred into 5 mL of sBC media in 1 well of a 6-well suspension plate to create clusters and placed in the 37°C incubator on an orbital shaker set at 95 rpm or seeded as single cells for sequential T cell assays.

### Dynamic Glucose Stimulated Insulin Secretion (GSIS) Assay

Dynamic insulin secretion was measured using a BioRep Technologies perifusion machine (PERI4-115-1810-076). 50 sBC clusters were placed on a filter in the perifusion chamber, and various solutions were perfused through the system at 100 μL/min by a peristaltic pump; cells and solutions were kept at 37°C and ambient atmosphere. The perifusion program consisted of a 30-minute preincubation with KRB buffer containing low (2.8 mM) glucose followed by the following run program: (i) 30 min low glucose, 20 min high (16.7 mM) glucose, 20 min high glucose + IBMX (50 μM, Millapore Sigma), 15 min low glucose, 5 min KCl (30 mM), 10 min low glucose. Perifusion flow-through was collected in 96 well plates and stored at 4°C overnight or - 20°C if longer storage was needed for future analysis. Cell pellets were recovered, lysed with acid/ethanol solution, and frozen overnight for assessment of total insulin content using a human insulin ELISA (ALPCO, 80-INSHU-E10.1).

### Immunofluorescence (IF) Imaging of sBC

Thawed sBC clusters were collected in 1.5 mL Eppendorf tubes and allowed to gravity settle for 5 min. Clusters were washed with 1x PBS and fixed for 10 min in 4% paraformaldehyde, blocked/permeabilized with CAS buffer containing 0.4% Triton X (CAS-T) for 10 min, and stained with primary antibodies overnight at 4°C in CAS-T buffer (**Table S2**). Clusters were washed and stained with fluorochrome-conjugated secondary antibodies for 2 hours at room temperature in CAS-T buffer, mounted on microscope slides (VWR) with ProLong Gold antifade reagent with DAPI (Invitrogen), and sealed with cover slips. Z-stack images were collected on Zeiss confocal microscope (LSM 710) followed by maximum intensity projection (Zeiss Black software).

### Human T Cell and NK Cell Subjects

Fresh peripheral blood mononuclear cells (PBMCs) were obtained from human leukapheresis-enriched blood of de-identified, IRB-exempt, healthy donors (T cell donors: median age: 24 years, range 19-43 years, *N*=11, 63.6% female; NK cell donors: median age: 33.5 years, range 18-49 years, *N*=4, 75% female) purchased from LifeSouth Community Blood Centers (Gainesville, FL, USA).

### Packaging and Titration of Lentiviral (LV) Vectors

The MART-1 (melanoma antigen-specific) T cell receptor (TCR) (37) and 1E6 (pre-proinsulin, PPI-specific)-TCR (38) LV constructs were packaged in third-generation lentivirus vectors containing a green-fluorescent protein (GFP) or Rat-CD2 reporter, respectively, produced by HEK293 cells co-transfected with the vesicular stomatitis virus G (VSV-G) envelope and Gag/Pol and Rev packaging plasmids, as previously described (39). 48 hours after transfection, viral supernatant was collected, centrifuged, and filtered before concentrating with PEG (VectorBuilder, Chicago, IL, USA). The infectivity of each vector was quantified as infectious units/mL (IFU/mL) by establishing the frequency of GFP^+^ or Rat-CD2^+^ cells compared to the volume of LV supernatant used. Subsequent transductions were performed at a concentration of 3 transducing units (TU)/cell where 1 IFU=1 TU.

### LV Transduction and Expansion of CD8^+^ T cells

Naïve CD8^+^ T cells were isolated from whole PBMCs using the EasySep™ Human Naïve CD8^+^ T Cell Isolation Kit II (StemCell Technologies, Vancouver, BC, Canada) and plated at 2.5 x 10^5^ cells/well in 1 mL of complete RPMI media (cRPMI; RPMI 1640 media Phenol Red w/o L-Glutamine 139 (Lonza, Basel, CH-BS, Switzerland), 5mM HEPES (Gibco, Waltham, MA, USA), 5 mM MEM 140 Non-Essential Amino Acids (NEAA; Gibco), 2mM Glutamax (Gibco), 50 µg/mL penicillin 141 (Gibco), 50 µg/mL streptomycin (Gibco), 20 mM sodium pyruvate (Gibco), 50 mM 2-mercaptoethanol (Sigma-Aldrich, St. Louis, MO, USA), 20 mM sodium hydroxide (Sigma-Aldrich) and 10% FBS (Genesee Scientific, El Cajon, CA, USA)). CD8^+^ T cells received 100 IU/mL rhIL-2 and were stimulated with Dynabeads™ Human T-Expander CD3/CD28 (Thermo Fisher Scientific, Waltham, MA, USA) at a 1:1 bead:cell ratio. After 48 hours of stimulation, cells were treated with protamine sulfate (8 µg/mL) and transduced with 3 TU/cell of the corresponding LV vector before spinnoculation (1000 g x 30 min at 32°C). Cell culture media was changed every 2-3 days, and beads were removed on day nine. To increase the purity of successfully transduced T cell “avatars” for co-culture experiments, fluorescence-activated cell sorting (FACS) was used to enrich GFP^+^ or Rat-CD2^+^ cells after expansion with a BD FACSMelody Cell Sorter (BD Biosciences, Franklin Lakes, NJ, USA).

### Human NK Cell Isolation

To verify that potential NK cell donors lacked the HLA-A2 serotype, 50,000 PBMCs were utilized for flow cytometry. Fc receptors were blocked using TruStain FcX (BioLegend, RRID: AB_2818986) for five min at 23°C before staining with an HLA-A2-Pacific Blue antibody (clone, RRID, concentration, and manufacturer information provided in **Table S2**) for 15 minutes at 23°C. Cells were washed once with stain buffer (PBS + 2% FBS + 0.05% NaN_3_ w/v) before analysis. Data were collected on an Aurora 5L (16UV-16V-14B-10YG-8R) spectral flow cytometer (Cytek, Freemont, CA, USA) and analyzed using FlowJo software (TreeStar; version 10.8.1). Using donors without the HLA-A2 serotype, human primary NK cells were isolated from whole PBMCs using the EasySep™ Human NK Isolation Kit (StemCell Technologies) for subsequent experiments.

### Immunoglobulin Binding Assay

Recombinant human TIGIT-Immunoglobulin (TIGIT-Ig; BioLegend, San Diego, CA, USA) and CD226-Immunoglobulin (CD226-Ig; BioLegfend) chimeric proteins were fluorescently labeled using a Zenon™ Alexa Fluor 594 (AF-594) human IgG labeling kit (Thermo Fisher Scientific) according to the manufacturer’s protocol, yielding a final concentration of 88 ng/µL of AF-594-labeled TIGIT-Ig and AF-594-labeled CD226-Ig. 50,000 cells were stained with Live/Dead™ Near-IR viability dye (Invitrogen, Waltham, MA, USA) for 10 min at 4°C before washing with stain buffer. Cells were resuspended at a concentration of 2.5 x 10^5^ cells/mL in 1x PBS (Gibco) and treated with AF-594-labeled TIGIT-Ig or AF-594-labeled CD226-Ig for 30 min at 37°C at the following concentrations: 0, 1, 10, 100, 50, 1000, 2500, 5000, 10000, and 25000 ng/mL. Unbound TIGIT-Ig or CD226-Ig was removed by washing cells with stain buffer. Flow cytometry data were collected and analyzed as described above.

### sBC Flow Cytometry Phenotyping

sBC were pre-treated with or without IFN-γ (100 ng/mL) for two days. 2 x 10^5^ cells from each condition were used for flow cytometry and were stained with Live/Dead™ Near-IR viability dye as described above. Cells were stained with an extracellular antibody cocktail consisting of anti-human CD112-PE/Cyanine7, CD155-Alexa Fluor 647, CD274-BV711, HLA-A2-Pacific Blue, and HLA-A,B,C-PE for 30 min at 23°C (clone, RRID, concentration, and manufacturer information provided in **Table S2**). Flow cytometry data were collected and analyzed as described above. Gating strategies were determined using fluorescence-minus one (FMO) and unstained controls.

### T Cell and NK Cell Activation Assay

To assess T cell or NK cell activation by flow cytometry after co-culture with sBC, 30,000 single sBC were plated on Cultrex-coated wells in a 96-well plate in day 16-23 media (described above) followed by 48 hours of IFN-γ treatment (100 ng/mL). sBC were then washed with complete DMEM media (cDMEM; Dulbecco’s Modification of Eagle’s Medium (DMEM; Lonza), 5 mM HEPES (Gibco), 5 mM MEM NEAA (Gibco), 50 µg/mL penicillin 141 (Gibco), 50 µg/mL streptomycin (Gibco), 0.02% Bovine Serum Albumin (BSA; Sigma-Aldrich), and 10% FBS (Genesee Scientific)) and co-cultured with MART-1 or PPI-reactive (1E6) T cell avatars at 10:1 effector:target (E:T) ratios for 48 hours or with primary NK cells at a 1:1 E:T ratio for 24 or 48 hours. After co-culture, immune cells were harvested and stained with Live/Dead™ Near-IR viability dye as described above. Following this, Fc receptors were blocked using TruStain FcX (BioLegend, RRID: AB_2818986) for five min at 23°C before extracellular staining in the presence of Brilliant Stain Buffer Plus (BD Biosciences) with panels for either T cell (CD8-AF700, CD96-BV421, CD226-PE-Cy7, TIGIT-PerCP eFluor710) or NK cell (CD3-AF700, CD27-Pacific Blue, CD56-SparkPLUS UV395, CD69-BV711, CD314-PE, HLA-DR-BV570) markers of interest for 30 min at 23°C (clone, RRID, concentration, and manufacturer information provided in **Table S2**). Flow cytometry data were collected and analyzed as described above. Gating strategies were determined using fluorescence-minus one (FMO) and unstained controls.

### Proliferation Assay

To assess the immunogenicity of each sBC line towards initiating an allogeneic, proliferative response in polyclonal naïve CD8^+^ T cells, we performed T cell proliferation assays. Briefly, naïve CD8+ T cells were isolated, as described above, and labeled with CellTrace™ Violet (CTV; Thermo Fisher, Cat: C34557A) proliferation tracking dye, as recommended by the manufacturer’s protocol. Subsequently, 187,500 labeled naïve CD8^+^ T cells were cultured at a 1:1 ratio with two-day IFN-γ pre-treated sBC for each condition in a 24-well plate and supplemented with rhIL-2 at 100 IU/mL. After six days of co-culture, suspension cells were collected and underwent viability staining with Live/Dead™ Near-IR viability dye and Fc receptor blocking, as described above before staining with CD8-AF700 for 30 min at 23°C (clone, RRID, concentration, and manufacturer information provided in **Table S2**). Flow cytometry data were collected and analyzed using proliferation modeling to determine the proliferation index using FlowJo software, as described above.

### Chromium-Release Assay

The susceptibility of sBC lines to cell-mediated lysis (CML) by T cell avatars and primary NK cells was assessed using chromium-release assays in the format previously described by Chen et al (40). sBC were plated and treated with IFNγ as described above before radiolabeling with ^51^CrNa_2_O_4_ (Revvity, Waltham, MA, USA) at an activity of 1.48 x 10^5^ Bq/well for four hours before washing twice with fresh cDMEM. sBC were co-cultured with MART-1 or PPI-reactive (1E6) T cell avatars at 0:1, 1:1, 5:1, and 10:1 effector:target (E:T) ratios for 48 hours or with primary human NK cells for 24 hours.

For chromium-release assays involving TIGIT blockade, PPI-reactive T cells or primary NK cells were concentrated at 3 x 10^6^ cells/mL cDMEM and incubated for 30 minutes at 37°C on an orbital shaker in the presence of 40 µg/mL of either mouse IgG2b, kappa isotype control (BioLegend; RRID: AB_2744505) or anti-human TIGIT mAb (BioLegend; RRID: AB_2820102) before washing into fresh cDMEM and plating as described above.

Following co-culture, the supernatants were removed, and along with 200 µL of 1x PBS used to wash the wells, were transferred into 6×50 mm lime glass tubes. The lysates of adherent cells were collected using a 2% SDS wash and transferred into separate tubes. ^51^Cr activity, measured in counts per minute (CPM), was assessed for both fractions on a Wizard 1470 automatic gamma counter (Revvity). The specific lysis of sBC was calculated as follows:

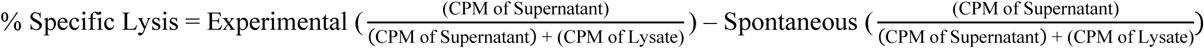

### Cytokine Multiplex Assay

To quantify the production of cytokines by T cells after co-culture with sBC, cell culture supernatants from the CML assay were used to evaluate IL-2, IL-4, IL-6, IL-10, IL-17A, FasL, IFN-γ, TNF, granzyme A, granzyme B, perforin, and granulysin production. Using the LEGENDplex™ Human CD8/NK Panel Kit (BioLegend), production of IFN-γ and granzyme A were measured at a 1:100 dilution, whereas FasL, granzyme B, perforin, and granulysin were measured at a 1:1 dilution, with IL-2, IL-4, IL-6, IL-10, IL-17A, and TNF not detected in the assay, using a 5L Aurora spectral flow cytometer. Data was analyzed using the LEGENDplex Data Analysis Software Suite (version 2024-09-10; BioLegend). Dilution factors and analyte detection ranges are described in **Table S3**.

### Data Visualization and Statistical Analysis

Statistical analyses were performed using GraphPad Prism software (version 10.3.1; San Diego, CA, USA). Unless otherwise stated, chromium-release and flow cytometric data were analyzed by two-way ANOVA, with Bonferroni’s post hoc test for multiple testing correction. Proliferation indices and binding area under the curve (AUC) values were analyzed using one-way ANOVA with Bonferroni’s post hoc test for multiple testing correction. P-values ≤ 0.05 were considered significant.

## Results

### Generation of WT and Mut CD155 hPSC Lines

To test whether CD155 expression can be utilized to reduce or prevent T cell activation by sBC, we introduced a genetically encoded expression cassette into the hPSC line Mel1^INS-GFP^, which contains a GFP reporter gene driven by the endogenous insulin promoter (pINS.GFP) (35). Using established protocols for TALEN-mediated site-specific genetic engineering of the AAVS1 locus in hPSC (19, 20), we targeted Mel1^INS-GFP^ hPSC to constitutively express WT or Mut CD155 in (**Figure 1A, Supp. Figure 1B**) and confirmed that targeting did not negatively impact the overt hPSC morphology of any isolated clonal hPSC line. Furthermore, protein expression of pluripotency markers OCT3/4 and NANOG was readily detectable by immunofluorescence (IF) staining (**Figure 1B**). Successful modification of clonal colonies was evaluated via flow cytometry to assess expression of CD155(**Figure 1C**).

**Fig. 1.**
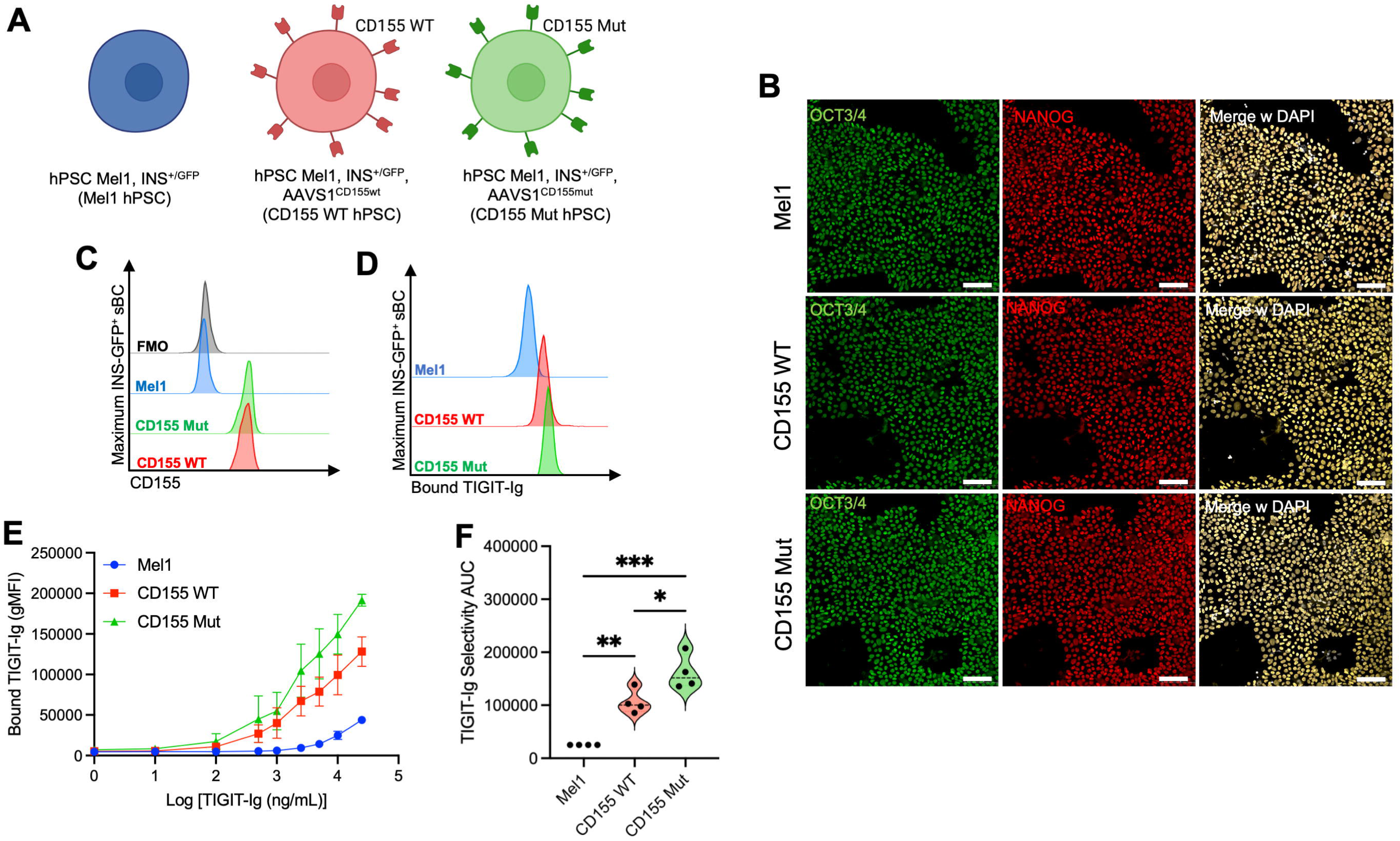
Genome engineering of CD155 WT and CD155 Mut Expressing hPSC. (**A**) Schematic of parental Mel1^INS-GFP^ (blue) and engineered WT (red) or Mut CD155 (green) hPSC lines. (**B**) Immunofluorescence staining for OCT3/4 (green), NANOG (red), and DAPI (white) in hPSC. (**C**) Flow cytometry for expression of CD155 in hPSC. (**D-F**) hPSC were treated with fluorescently labeled TIGIT-Ig to compare binding efficiency between the parental Mel1 (blue), CD155 WT-expressing (red), and CD155 Mut-expressing (green) lines. (**D**) Representative histograms of cell-bound Ig after treatment at the 25,000 ng/mL condition. (**E**) Differences in Ig binding efficiency characterized across a range of 0-25,000 ng/mL, with additional comparisons made using (**F**) TIGIT-Ig selectivity area under the curve (AUC) values for each binding curve, where the bound TIGIT-Ig gMFI value for each line was normalized to the Mel1-bound TIGIT-Ig gMFI at each concentration. Data reflects biological n=3/condition for TIGIT-Ig sBC. Significant P-values are reported for one-way ANOVA with Bonferroni’s multiple comparisons of AUC values between paired samples. *p<0.05, **p<0.01, ***p<0.001.

To characterize if the CD155 WT or Mut expression influenced binding of the ligand TIGIT, we cultured hPSC with fluorescently labeled TIGIT-Ig fusion protein. Across a range of 0-25,000 ng/mL, we observed that both CD155 WT and CD155 Mut hPSC displayed greater binding efficiency for TIGIT-Ig (**Figure 1D-F**; CD155 WT: 4.25-fold, p=0.0054; CD155 Mut:6.47-fold, p=0.0003) compared to the parental Mel1 hPSC. Notably, the CD155 Mut line displayed greater cell binding for TIGIT-Ig (1.52-fold, p=0.033) compared to the CD155 WT line. These differences were not observed with a non-specific isotype control Ig antibody (**Supplemental Figure 2A**).

### CD155 Expressing hPSC Efficiently Differentiate into sBC

Using our previously established methods (41), we then performed directed differentiation into sBC experiments with both modified hPSC lines, as well as the unmodified parental hPSC line as a control (**Figure 2A**). Flow cytometric analysis for pluripotency markers TRA-1-60 and SOX2 demonstrate uniform expression of pluripotency markers in all three hPSC lines at the start of differentiation, which was lost after DE generation at day three, supporting the IF data. As expected, DE markers FOXA2 and SOX17 were absent at the start but showed high expression by the third day of differentiation (**Supplemental Figure 1G-J**). At subsequent days of differentiation, the morphology of Mel1, CD155 WT, and CD155 Mut hPSC clusters displayed similar morphology and expression of the pINS.GFP reporter at the end of differentiation (**Figure 2B**). At day 23 of the protocol all hPSC lines reproducibly generated approximately ∼50% insulin-expressing sBC. We did not observe significant differences in the frequency of β-cell markers C-peptide (**Figure 2C**) and NKX6.1 (**Figure 2D**), or the frequency of alpha cell marker glucagon (**Figure 2E**) by flow cytometry. However, when we measured the expression of CD155 on sBC, we observed the retention of CD155 expression in virtually all the modified cells whether assessing total population (**Figure 2F**) or in C-peptide^+^ sBC cells (**Figure 2G**). The frequencies as well as gMFI of CD155 expression were significantly higher in edited sBC compared to Mel1 sBC (**Figure 2H**), similar to our hPSC analysis for transgene expression. IF staining of day 23 sBC revealed comparable morphology and expression of insulin and NKX6.1 across all three lines (**Figure 2I**). To assess sBC function, we conducted dynamic GSIS assays and demonstrated that CD155 WT and CD155 Mut sBC are similarly responsive to high glucose, IBMX (triggering the amplifying pathway) (42), and depolarization with KCl compared to Mel1 sBC controls (**Figure 2J**). Collectively, these data demonstrate that expression of CD155 does not influence the differentiation efficiency or function of sBC.

**Fig. 2.**
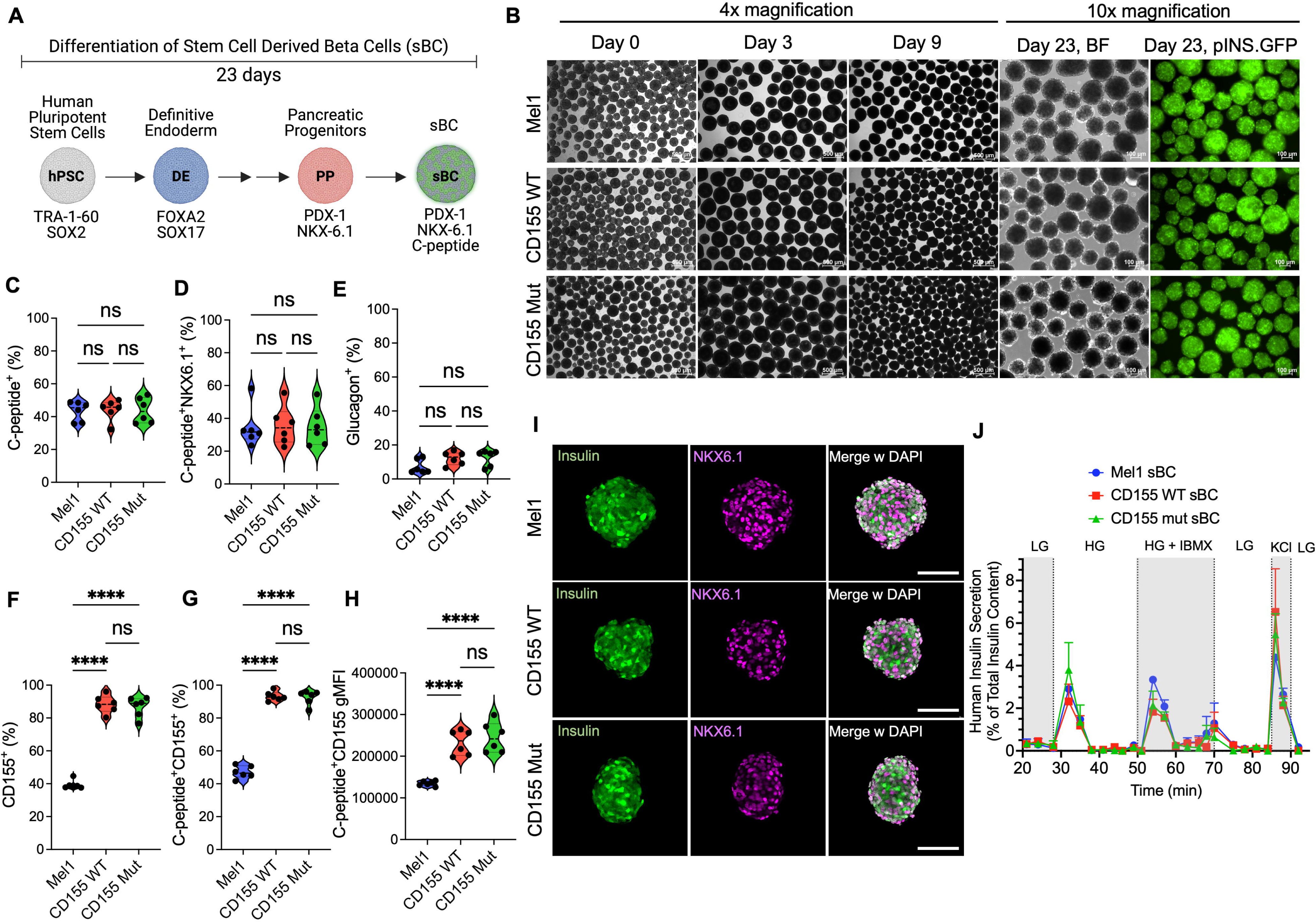
CD155 WT and CD155 Mut sBC show similar differentiation and function. (**A**) Schematic of differentiation from hPSC to sBC. (**B**) Representative bright field and pINS.GFP reporter live images of differentiating clusters at subsequent time points. 4x or 10x magnification images as denoted. (**C-F**) Flow cytometry analysis at day 23 of n=6 independent differentiations of Mel1, CD155 WT, and CD155 Mutant sBC for frequency of (**C**) C-peptide^+^ cells, (**D**) C-peptide^+^ NKX6.1^+^ double positive cells, (**E**) glucagon^+^ cells, (**F**) total CD155^+^ cells or (**G**) CD155^+^ C-peptide^+^ populations, and (**H**) geometric mean fluorescence intensity (gMFI) of CD155 within C-peptide^+^ cells. (**I**) Immunofluorescence staining of day 23 sBC for insulin (green), NKX6.1 (magenta), and DAPI (white). (**J**) Dynamic glucose stimulated insulin secretion data from perfusion study as percent of total insulin content. n=3 independent differentiations. Significant P-values are reported for one-way ANOVA with Bonferroni’s multiple comparisons. ****p<0.0001.

### Mutant CD155 Exhibits Greater Binding for TIGIT-Ig and CD226-Ig

To characterize how the expression of either WT or Mut CD155 on sBC impacted binding towards TIGIT or CD226, we cultured sBC with fluorescently labeled TIGIT-Ig fusion protein. Across a range of 0-25,000 ng/mL, we observed that both CD155 WT and CD155 Mut sBC displayed greater binding efficiency for TIGIT-Ig (**Figure 3A-C**; CD155 WT: 2.14-fold, p=0.0064; CD155 Mut:2.93-fold, p=0.0008) and CD226-Ig (**Figure 3D-F**; CD155 WT: 1.33-fold, p=0.0013; CD155 Mut: 1.73-fold, p<0.0001) compared to the parental Mel1 sBC. Notably, the CD155 Mut line displayed greater binding for both TIGIT-Ig (1.37-fold, p=0.024) and CD226-Ig (1.29-fold, p=0.0005) as compared to the CD155 WT line. Using a control isotype Ig in the same assay demonstrated no significant differences between the three lines (**Supplemental Figure 2B**). These data support previously reported biolayer interferometry (BLI) results by Matsuo *et al.*, suggesting that Mut CD155 demonstrates a greater efficiency for the ligands TIGIT and CD226 (34).

**Fig. 3.**
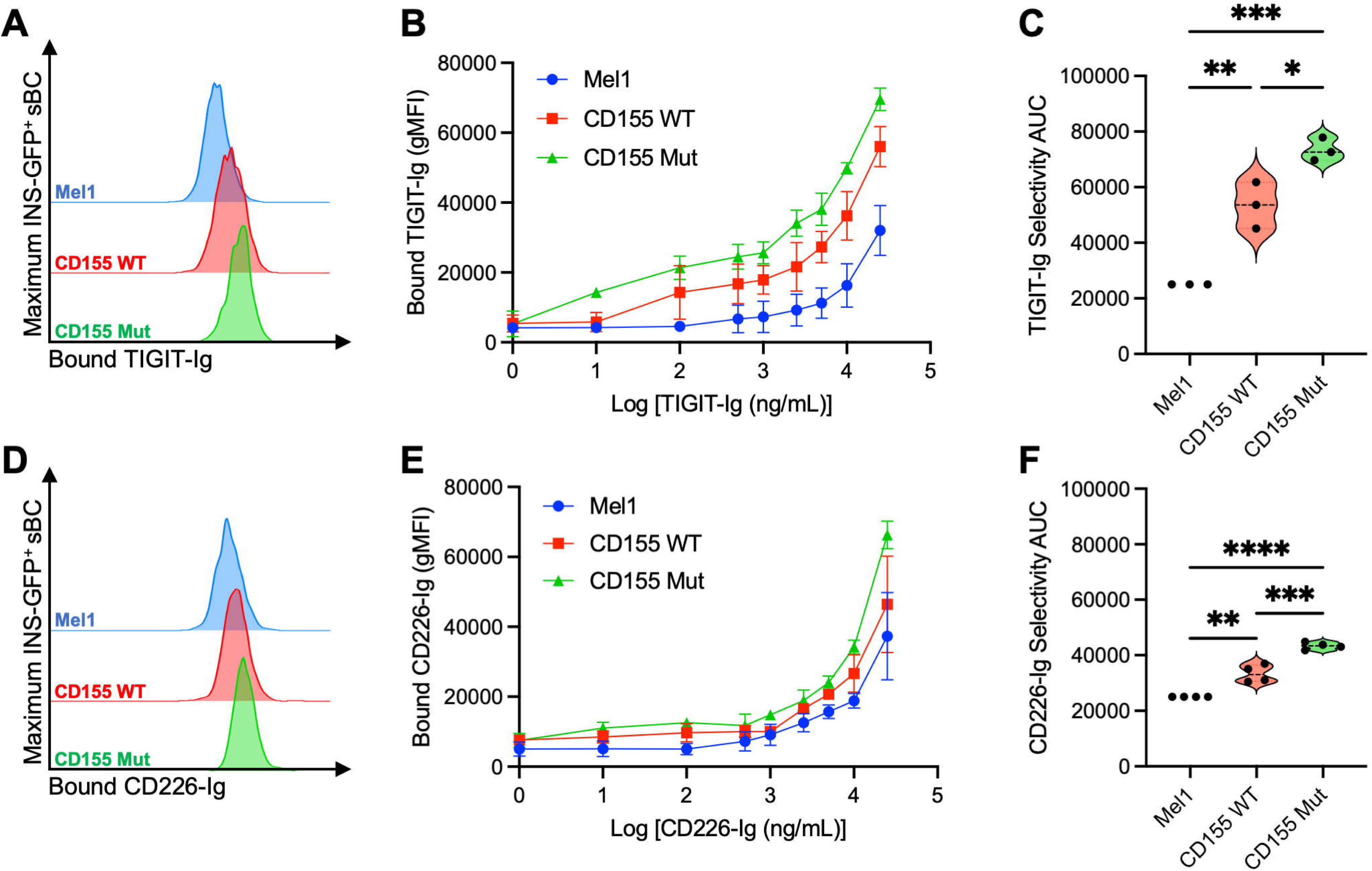
Expression of Mutant CD155 Confers Greater Binding for TIGIT-Ig and CD226-Ig on sBC. sBC were treated with fluorescently labeled (**A-C**) TIGIT-Ig or (**D-F**) CD226-Ig to compare binding efficiency between the parental Mel1 (blue), CD155 WT-expressing (red), and CD155 Mut-expressing (green) lines. (**A, D**) Representative histograms shows cell-bound Ig after treatment at the 25,000 ng/mL condition. (**B, E**) Differences in Ig binding efficiency were characterized across a range of 0-25,000 ng/mL, with additional comparisons made using (**C, F**) area under the curve (AUC) values for each binding curve. Data reflects biological n=3/condition for TIGIT-Ig sBC and n=4/condition for CD226-Ig. Significant P-values are reported for one-way ANOVA with Bonferroni’s multiple comparisons of AUC values between paired samples. *p<0.05, **p<0.01, ***p<0.001. ****p<0.0001.

### IFN-γ Does Not Upregulate CD112 or CD155 Expression on sBC

In the inflamed pancreatic islet microenvironment during the pathogenesis of T1D, IFN-γ has been shown to promote immunogenicity through upregulating MHC-I molecules (43, 44). Indeed, we observed greater expression of HLA-A2 (**Figure 4A-B**) and HLA-A,B,C (**Figure 4C-D**) following IFN-γ treatment across all three sBC lines, as well as upregulation of PD-L1 (**Figure 4E-F**) as we have previously reported (19). These data indicate that CD155 genome engineering does not negate prototypical mechanisms of standard immune surveillance or response to cytokine treatment, which would be a caveat of any immunological assessments. However, we did observe reductions in CD112 (**Figure 4G-H**) expression, another ligand that can bind TIGIT and CD226 (45), on CD155 expressing sBC compared to Mel1 controls as well as small but significant reductions in HLA-A,B,C and PD-L1 in IFNγ-treated conditions. The very high levels of CD155 expression driven by the CAG promoter in our system may be limiting available cellular surface receptor space causing reductions in some receptor gMFI upon IFNγ stimulation (46). However, as modified sBC are still capable of upregulating non-CD155 surface receptor expression upon IFNγ stimulation, the observed small differences may not have biological significance. Nonetheless, we did not detect a significant impact on the expression of CD112 or CD155 (**Figure 4I-J**) after IFNγ treatment, suggesting the expression of these ligands are not impacted by the cytokine treatment.

**Fig. 4.**
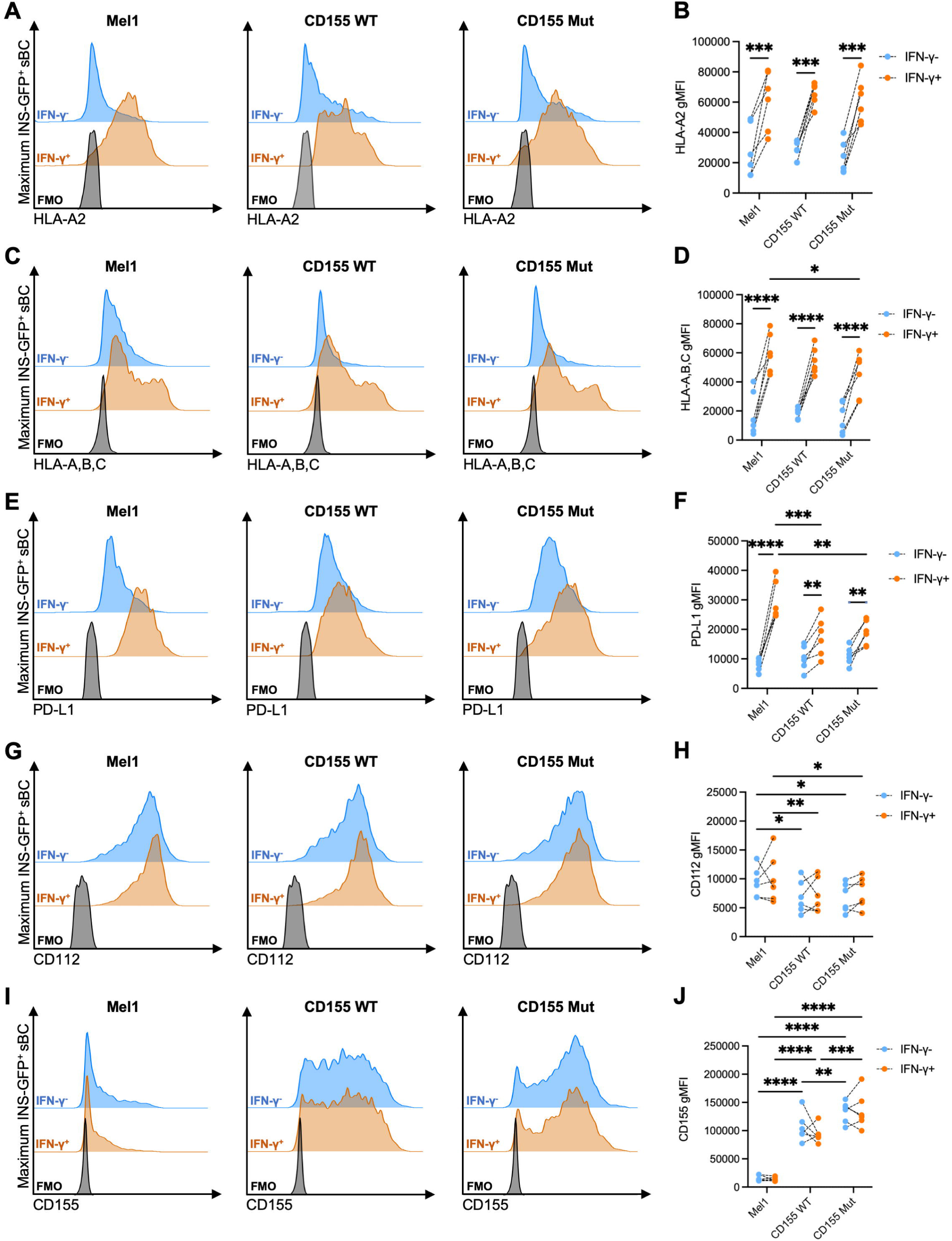
IFN-γ Treatment Does Not Upregulate CD112 or CD155 Expression on sBC. Expression of (**A, B**) HLA-A2, (**C, D**) HLA-A,B,C, (**E, F**) PD-L1, (**G, H**) CD112, and (**I, J**) CD155 was evaluated on INS-GFP^+^ sBC following 48 hours with (orange) or without (blue) IFN-γ treatment. (**A, C, E, G, I**) Representative histograms show staining of Mel1, CD155-WT, and CD155-Mut sBC lines relative to FMO controls (black). (**B, D, F, H, J**) Paired dot plots show gMFI values by sBC line and treatment condition. Data reflects biological n=6/condition. Significant P-values are reported for two-way ANOVA with Bonferroni’s multiple comparisons between paired samples. *p<0.05, **p<0.01, ***p<0.001, ****p<0.0001.

### CD8^+^ T cells Demonstrate Reduced Activation Following Co-Culture with CD155 Mut sBC

To understand whether expression of the WT or Mut CD155 on sBC impacts the activation of responder CD8^+^ T cells, we performed HLA-peptide-TCR matched co-culture assays with T cell avatars specific for an irrelevant antigen (clone MART-1) or towards PPI (clone 1E6), a canonical autoantigen presented to T cells by sBC at an E:T ratio of 10:1 (**Figure 5A**). Following 48 hours of co-culture with CD155 Mut sBC, when examining the expression of the CD155 ligands, we observed that MART-1 and PPI-reactive avatars co-cultured with CD155 Mut sBC displayed reduced expression of CD96 (**Figure 5B-E**), compared to avatars co-cultured with CD155 WT (MART-1: 0.69-fold, p=0.0025; PPI: 0.74-fold, p=0.031) or Mel1 (MART-1: 0.69-fold, p=0.0019; PPI: 0.74-fold, p=0.031) sBC with even more apparent reductions in CD226 expression (**Figure 5F-I**) compared to avatars co-cultured with CD155 WT (MART-1: 0.46-fold, p=0.0024; PPI: 0.50-fold, p=0.0097) or Mel1 (MART-1: 0.31-fold, p<0.0001; PPI: 0.40-fold, p=0.0004) sBC. Interestingly, MART-1 T cell avatars demonstrated similar levels of TIGIT expression after co-culture with the different sBC (**Figure 5J-K**), while PPI-reactive avatars displayed a significant reduction in TIGIT expression after co-culture with CD155-engineered sBC (**Figure 5L-M**), suggesting a potential antigen-specific difference in this pathway. These data suggest that the expression of mutant CD155 by sBC reduces the activation and functionality of CD8^+^ T cells following engagement.

**Fig. 5.**
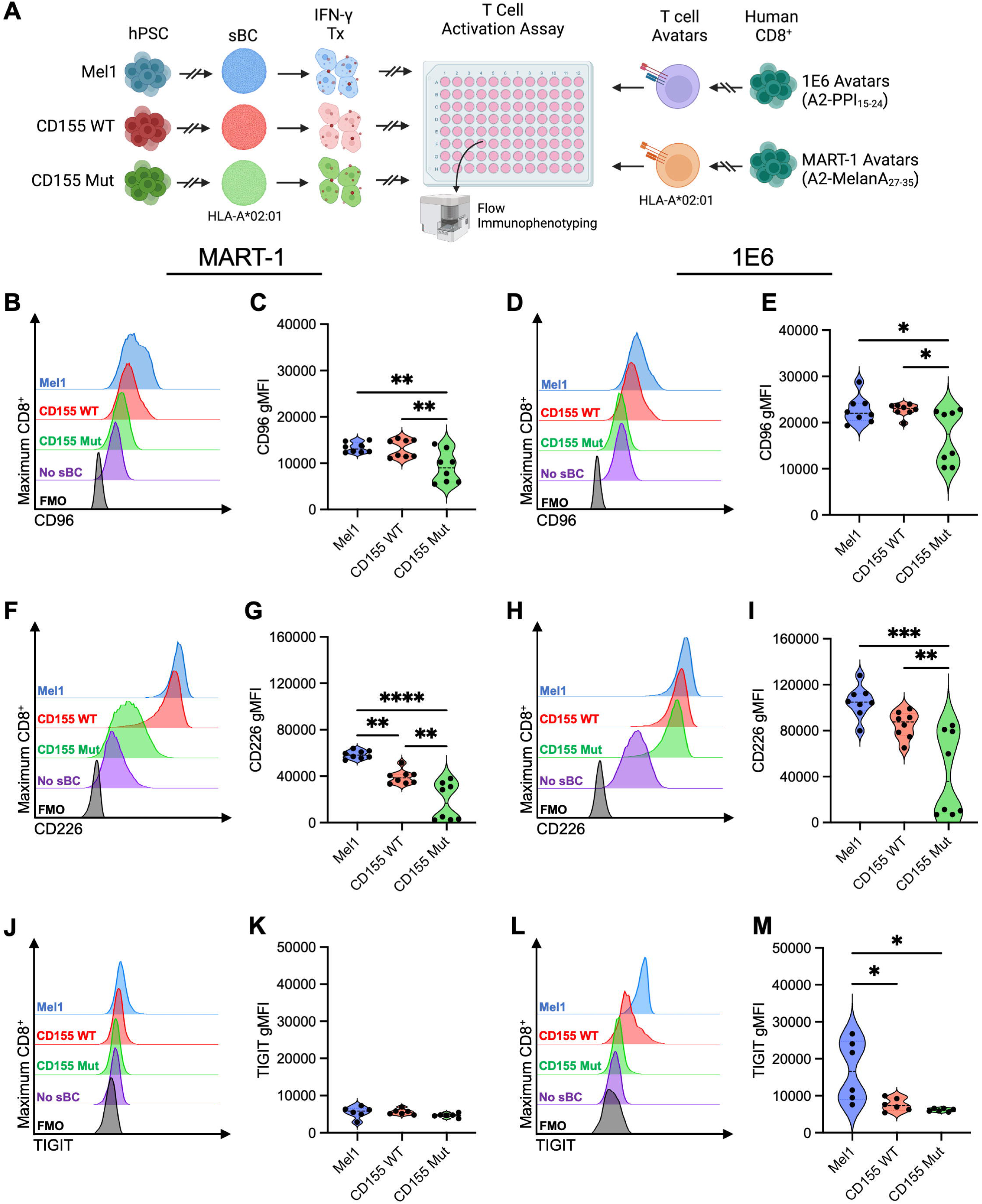
Expression of Mutant CD155 Reduces T cell Activation During HLA-peptide-TCR matched Co-culture. (**A**) Experimental scheme depicting activation assay to assess the immunogenicity of Mel1 (blue), CD155 WT-expressing (red), and CD155 Mut-expressing (green) sBC by MART or PPI-reactive avatars, as measured by flow immunophenotyping of T cells (created with BioRender). Plots show differences in expression of the T cell activation markers, (**B-E**) CD96, (**F-I**) CD226, and (**J-M**) TIGIT on (**B-C, F-G, J-K**) MART-1 or (**D-E, H-I, L-M**) PPI-reactive avatars between sBC lines at the 10:1 Effector:Target (E:T) ratio compared to no sBC (purple) and no dye (black) controls. Data reflects biological n=8/condition. Significant P-values are reported for one-way ANOVA with Bonferroni correction for multiple comparisons. *p<0.05, **p<0.01, ***p<0.001, ****p<0.0001.

To determine how the expression of Mut CD155 impacts allogeneic T cell responses, we cultured proliferation-dye labeled naïve CD8^+^ T cells with sBC to assess proliferative capacity (**Figure 6A**). Following six days of co-culture, we observed reductions in CD8^+^ T cell proliferation when cultured with CD155 WT (0.92-fold, p=0.0003) or CD155 Mut (0.84-fold, p<0.0001) sBC compared to the parental Mel1 sBC (**Figure 6B-C**). Furthermore, we identified greater reductions in CD8^+^ T cell proliferation in the CD155 Mut co-culture (0.92-fold, p=0.0006) compared to the CD155 WT co-culture (**Figure 6B-C**). Together, these data suggest that expression of CD155, particularly the Mut CD155, by sBC reduces immunogenicity from alloreactive T cells.

**Fig. 6.**
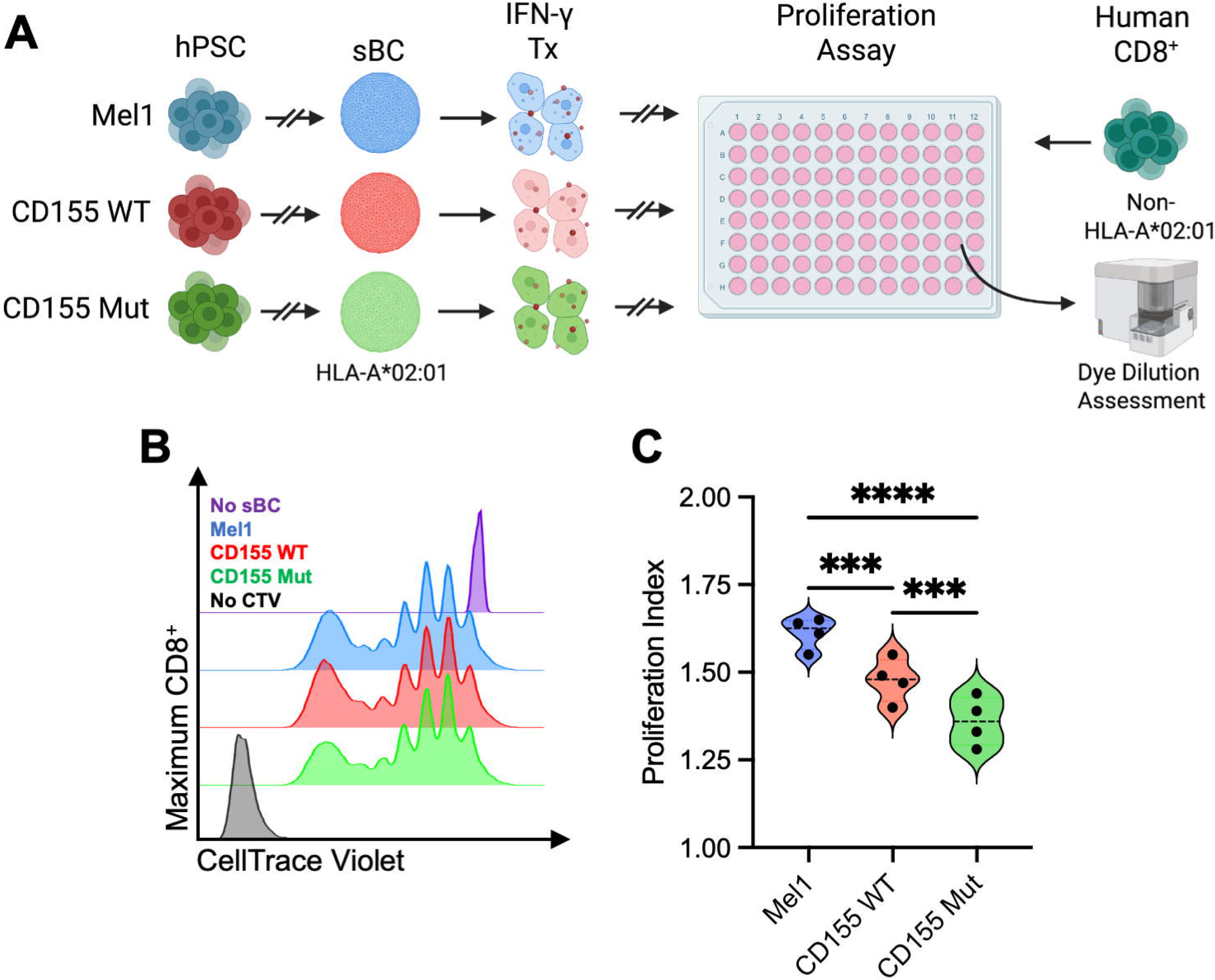
Expression of Mutant CD155 Reduces CD8^+^ T cell Allogeneic Proliferative Responses. (**A**) Experimental scheme depicting proliferation assay used to characterize the immunogenicity of sBC lines when cultured with allogenic naïve CD8^+^ T cells (created with BioRender). (**B**) Representative dye dilution plots depict the proliferation of naïve CD8^+^ T cells following six days of co-culture with Mel1 (blue), CD155 WT-expressing (red), and CD155 Mut-expressing (green) sBC compared to no sBC (purple) and no dye (black) controls, with (**C**) violin plots showing proliferation indices for each condition (biological n=4/condition). Significant P-values are reported for paired samples using one-way ANOVA, with Bonferroni correction for multiple comparisons. ***p<0.001, ****p<0.0001.

### Expression of CD155 Mut Confers Greater Resistance to sBC from T Cell-Mediated Lysis

To understand how the expression of either the WT or Mut CD155 impacts the cytotoxicity of T cells towards sBC, we performed co-culture assays with MART-1 and PPI avatars (**Figure 7A**) at varying E:T ratios of: 1:1, 1:5, and 1:10. After two days of co-culture with MART-1 avatars, we observed no significant changes in the cytolytic molecules FasL or granzyme B production between the CD155 WT and the CD155 Mut sBC lines (**Figure 7B-C**). However, at the 10:1 ratio, we observed that MART-1 avatars co-cultured with CD155 Mut sBC produced significantly less perforin (0.85-fold, p=0.034) and granulysin (0.74-fold, p=0.032) than avatars co-cultured with CD155 WT sBC (**Figure 7D-E**), suggesting that expression of Mut CD155 can reduce T cell cytotoxicity without antigen recognition. Further, in the context of antigen-specific recognition, we observed that PPI-reactive avatars cultured with CD155 Mut sBC at a 10:1 ratio produced significantly less FasL (0.52-fold, p<0.0001), granzyme B (0.56-fold, p=0.0003), perforin (0.66-fold, p<0.0001), and granulysin (0.75-fold, p=0.0013) compared to PPI-reactive avatars co-cultured with CD155 WT sBC (**Figure 7F-I**). Notably, we also observed significant decreases in the production of FasL (0.44-fold, p<0.0001) and perforin (0.91-fold, p=0.037) by PPI-reactive avatars cultured with CD155 Mut sBC at the 5:1 E:T ratio, compared to co-cultures with CD155 WT sBC.

**Fig. 7.**
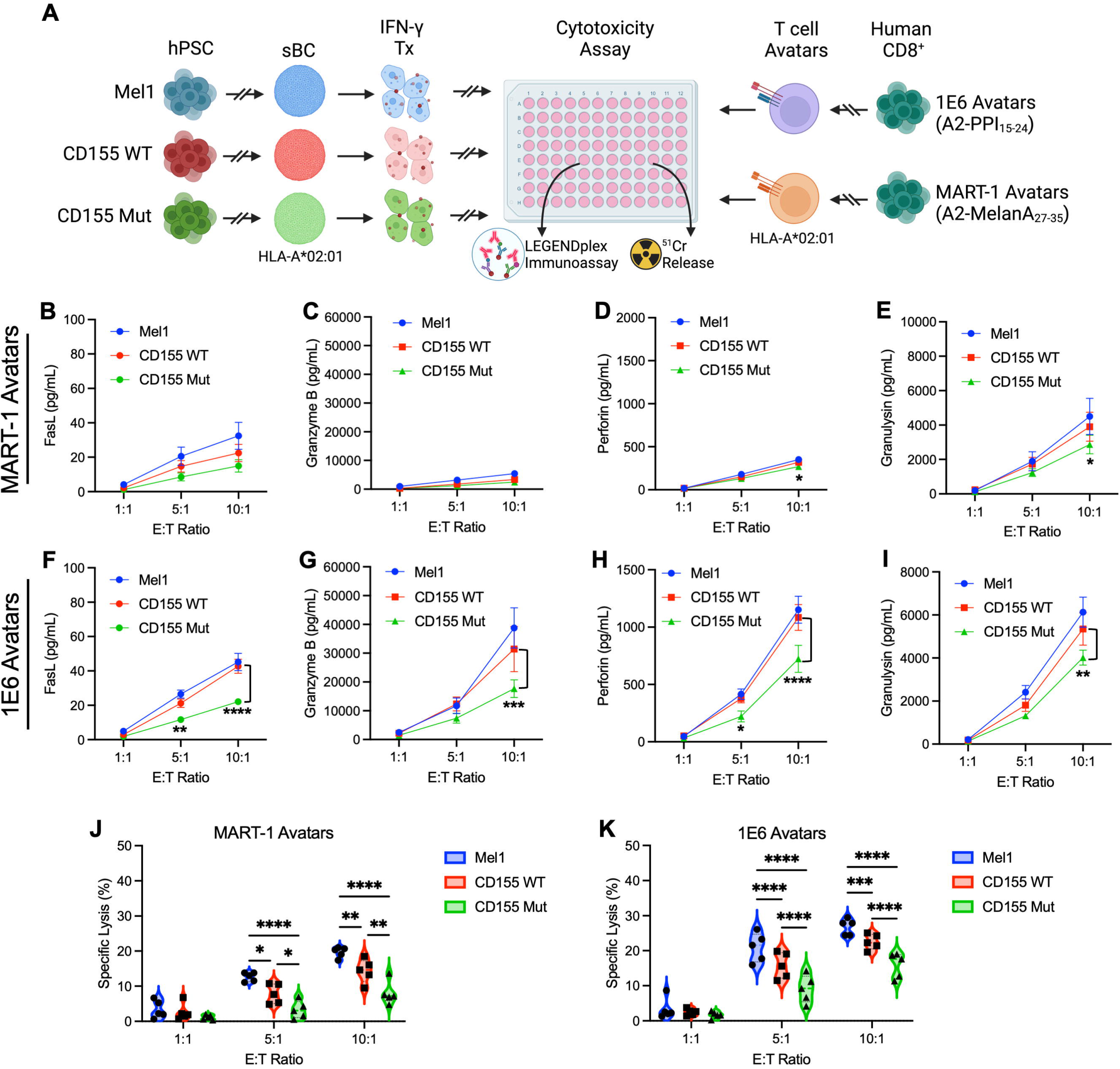
Expression of Mutant CD155 Provides sBC Greater Protection from T Cell-Mediated Lysis. (**A**) Experimental scheme depicting CML assay to assess the immunogenicity of Mel1 (blue), CD155 WT-expressing (red), and CD155 Mut-expressing (green) sBC by MART or PPI-reactive avatars, as measured by ^51^Cr-release, with T cell cytokine production assessed by LEGENDplex (created with BioRender). Plots show differences in cell culture supernatant concentrations following co-culture with (**B-E**) MART-1 or (**F-I**) PPI-reactive avatars of (**B, F**) FasL, (**C, G**) granzyme B, (**D, H**) perforin, and (**E, I**) granulysin between sBC lines at each Effector:Target (E:T) ratio. Data reflects biological n=8/condition. Significant P-values are reported for two-way ANOVA with Bonferroni correction for multiple comparisons between CD155 WT and CD155 Mut conditions. (**J, K**) Violin plots show percent-specific lysis of sBC by (**J**) MART-1 or (**K**) PPI-reactive avatars at each E:T ratio. Data reflects biological n=5/condition. Significant P-values are reported for two-way ANOVA with Bonferroni correction for multiple comparisons. *p<0.05, **p<0.01, ***p<0.001, ****p<0.0001.

To determine if these decreases in immunogenicity between the CD155 WT and CD155 Mut sBC lines translate to protection from CML, we assessed sBC killing by T cell avatars via chromium release assay. After 48 hours of co-culture, we identified that CD155 Mut sBC showed reductions in specific lysis at the 5:1 (0.42-fold, p=0.021) and 10:1 (0.56-fold, p=0.0015) E:T ratios following co-culture with MART-1 avatars, as compared to CD155 WT sBC (**Figure 7J**). Strikingly, we observed that CD155 Mut sBC also demonstrated increased protection from antigen-specific CML by PPI-reactive avatars, with significantly diminished lysis at the 5:1 (0.58-fold, p<0.0001) and 10:1 (0.71-fold, p<0.0001) E:T ratios compared to CD155 WT sBC (**Figure 7K**).

### TIGIT Blockade Ablates Protection Conferred by CD155 Expression on sBC

Given the increased protection from antigen-specific CML conferred to sBC with increased availability and higher affinity CD155 Mut, we sought to determine the mechanism accounting for the reduced cytotoxic activity of CD8^+^ T cell avatars. To accomplish this, we evaluated the contribution of CD155:TIGIT signaling by treating PPI-reactive avatar T cells with an anti-TIGIT blocking antibody before co-culture with Mel1 or CD155 Mut sBC (**Figure 8A**). Compared to an isotype control, we did not observe significant differences in T cell cytolytic molecule production resulting from TIGIT blockade when PPI-reactive avatars were co-cultured with Mel1 sBC (**Figure 8B-E, grey lines**), which would not be expected due to low CD155:TIGIT signaling. However, when TIGIT was blocked on PPI-reactive avatars and co-cultured with CD155 Mut sBC (**Figure 8B-E, pink lines**), we observed greater production of the cytolytic molecules FasL (**Figure 8B**; 10:1: 1.71-fold, p=0.0035), granzyme B (**Figure 8C**; 10:1: 2.43-fold, p=0.017), perforin (**Figure 8D**; 10:1: 3.06-fold, p<0.0001), and granulysin (**Figure 8E**; 10:1: 1.97-fold, p=0.0002), compared to PPI-reactive avatars treated with an isotype control.

**Fig. 8.**
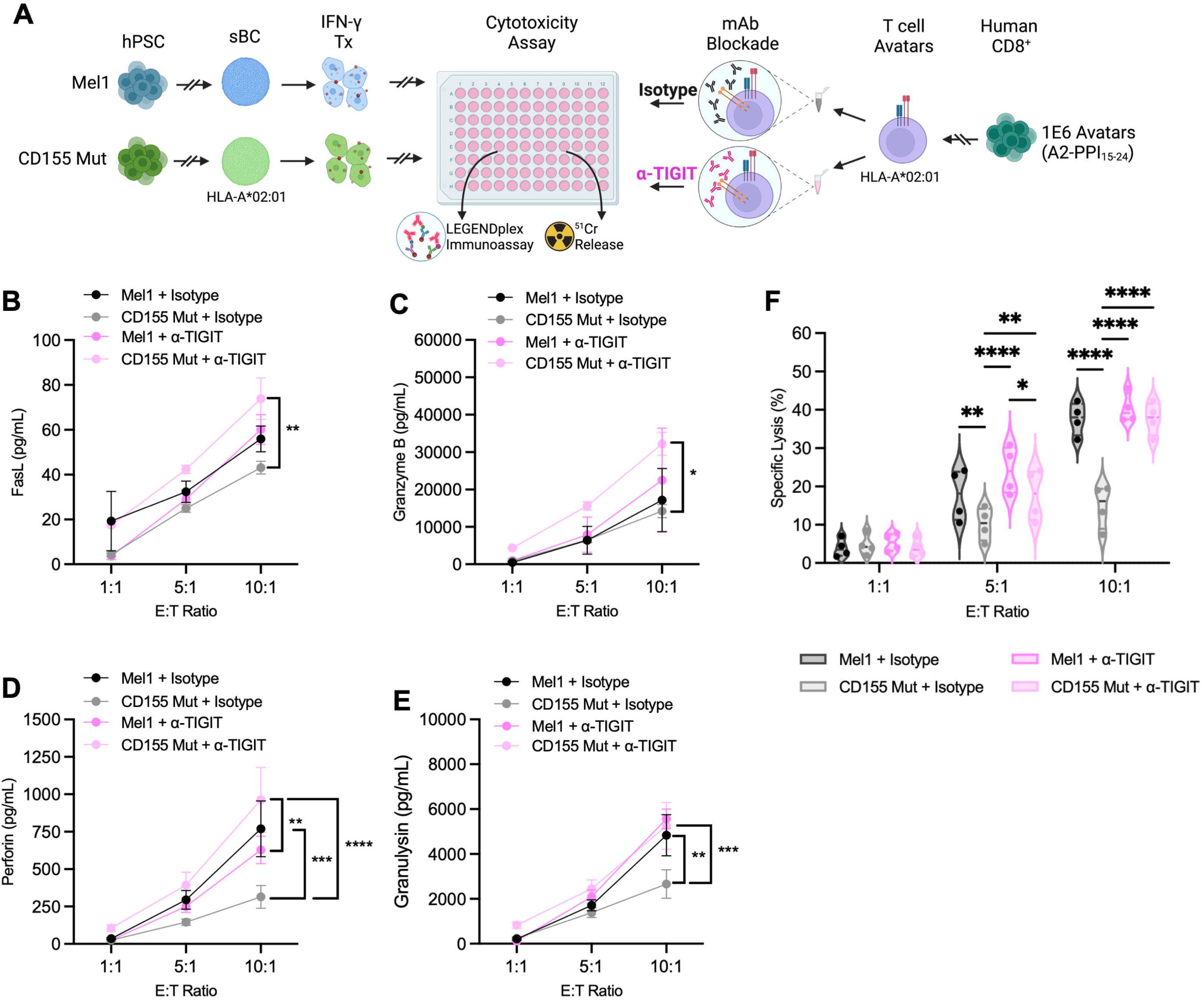
TIGIT Blockade Ablates sBC Protection from CML Conferred by Mut CD155 Expression. (**A**) Experimental scheme depicting CML assay to assess the effect of TIGIT blockade (pink), relative to an isotype control (black) on CML of parental Mel1 and CD155 Mut-expressing sBC lines by PPI-reactive avatars, as measured by ^51^Cr-release, with T cell cytokine production assessed by LEGENDplex (created with BioRender). Plots show differences in cell culture supernatant concentrations following co-culture with Mel1 or CD155 Mut sBC of (**B**) FasL, (**C**) granzyme B, (**D**) perforin, and (**E**) granulysin between ⍺-TIGIT and isotype treated PPI-reactive avatars at each Effector:Target (E:T) ratio. Data reflects biological n=4/condition. (**F**) Violin plots show percent-specific lysis of Mel1 (dark grey and dark pink) or CD155 Mut (light grey and light pink) sBC at each E:T ratio with or without α-TIGIT blockade. Data reflects biological n=4/condition. Significant P-values are reported for three-way ANOVA with Bonferroni correction for multiple comparisons. *p<0.05, **p<0.01, ***p<0.001, ****p<0.0001.

When assessing sBC killing by chromium-release assays, we again observed a protective impact of CD155 Mut expression on sBC compared to Mel1 control sBC cultures in the presence of the Isotype antibody blockade (**Figure 8J, grey bars**). Upon TIGIT blockade, we did not see any significant changes in the susceptibility of Mel1 sBC to CML (**Figure 8F, dark grey to dark pink bars**), but we observed significantly increased killing of CD155 Mut sBC to CML as a result of TIGIT blockade at both the 5:1 (1.80-fold, p=0.0058) and 10:1 (1.91-fold, p<0.0001) E:T ratios (**Figure 8F, light grey to light pink bars**). In fact, TIGIT blockade returned CD155 Mut sBC CML levels to those equivalent to Mel1 control sBC co-cultures. Together, these data suggest that the increased protection conferred to sBC from antigen-specific CML by the expression of Mut CD155 is a result of more robust coinhibitory CD155:TIGIT signaling, the loss of which likely skews T cells toward preferential CD226:CD155 costimulation and augmented cytotoxic potential.

### CD155 Expression in sBC Suppresses NK Activation and NK-mediated Cytotoxicity

In addition to T cell responses, innate immune recognition of transplanted beta cells, especially in an allogeneic transplant context, can facilitate graft destruction. One innate immune subset that utilizes TIGIT signaling similarly to T cells is natural killer (NK) cells (47). As a result, we interrogated the ability of CD155 expressing sBC to inhibit NK activation and cell-mediated killing. HLA-mismatched NK cells were isolated from PBMC donors followed by incubation with IFNγ treated Mel1, CD155 WT, or CD155 Mut sBC (**Figure 9A**). Flow cytometry analysis of the NK cell activation markers CD27 (**Figure 9B, 9C**), CD69 (**Figure 9D, 9E**), NKG2D (**Figure 9F, 9G**), and HLA-DR (**Figure 9H, 9I**) displayed significant reductions in receptor expression gMFI after co-culture within CD155 Mut sBC co-cultures (CD27: 0.83-fold, p<0.0001; CD69: 0.77-fold, p=0.0013; NKG2D: 0.70-fold, p<0.0001; HLA-DR: 0.79-fold, p<0.0001) compared to Mel1 controls (48–51). Like in T cell assays, CD155 WT expressing sBC displayed a more intermediate suppressive impact, with significant reductions in NKG2D (0.72-fold, p<0.0001) compared to Mel1 controls, while the other receptors’ expression was not significantly altered.

**Fig. 9.**
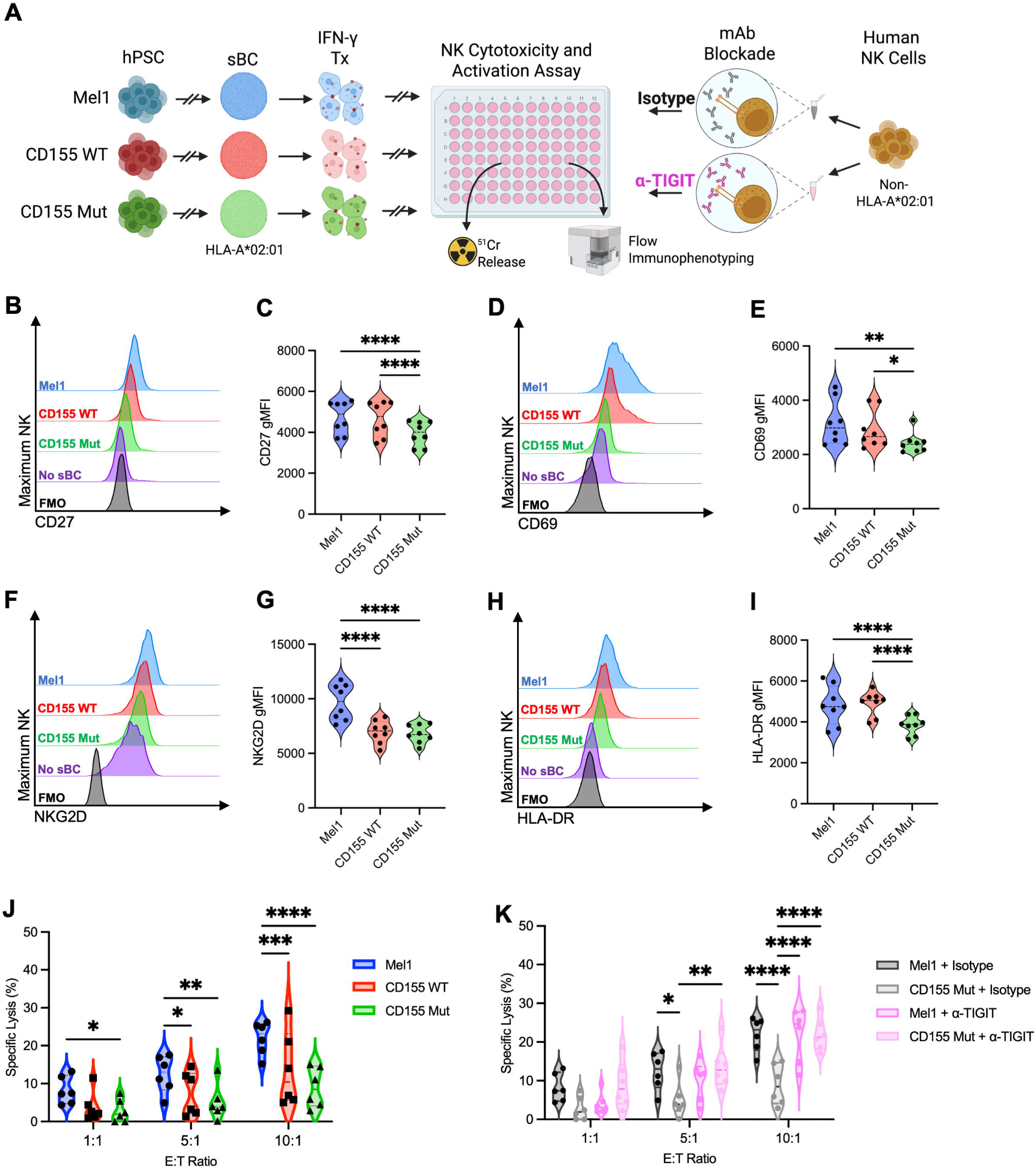
CD155 Expression in sBC Suppresses NK Activation and NK-mediated Cytotoxicity. **(A)** Experimental scheme depicting CML assay to assess the effect of TIGIT blockade (pink), relative to an isotype control (black) on CML of Mel1 (blue), CD155 WT-expressing (red), and CD155 Mut-expressing (green) sBC by NK cells, as measured by flow immunophenotyping of NK cells and ^51^Cr-release assays (created with BioRender). (**B-I**) Violin plots show differences in expression of the NK cell activation markers (**B, C**) CD27, (**D, E**) CD69, (**F, G**) NKG2D, and (**H, I**) HLA-DR between sBC lines at the 1:1 Effector:Target (E:T) ratio compared to no sBC (purple) and no dye (black) controls. Data reflects biological n=8/condition. Significant P-values are reported for one-way ANOVA with Bonferroni correction for multiple comparisons. (**J, K**) Violin plots show percent-specific lysis of (**J**) Mel1, CD155 WT, and CD155 Mut sBC at each E:T ratio in isotype control cultures and of (**K**) Mel1 (dark grey and dark pink) or CD155 Mut (light grey and light pink) sBC at each E:T ratio with or without α-TIGIT blockade. Data reflects biological n=6/condition. Significant P-values are reported for three-way ANOVA with Bonferroni correction for multiple comparisons. *p<0.05, **p<0.01, ***p<0.001, ****p<0.0001.

To determine the mechanistic role of CD155 expression on sBC within the NK cell assays, we pre-incubated NK cells with either an isotype control antibody or anti-TIGIT blocking antibody as in the T cell assays (**Figure 8**). Chromium release assays quantifying sBC lysis after co-culture experiments with the isotype control antibody revealed significant reductions in NK cell-mediated sBC lysis with CD155 Mut co-cultures compared to Mel1 controls (1:1: 0.34-fold, p=0.033; 5:1: 0.41-fold, p=0.0029; 10:1: 0.41-fold, p<0.0001) at all three effector:target ratios (**Figure 9J**). There were also significant reductions in specific lysis within CD155 WT sBC co-cultures (5:1 0.49-fold, p=0.047; 10:1: 0.62-fold, p=0.0010) at the 5:1 and 10:1 ratios compared to Mel1 controls. Upon blockade of CD155:TIGIT engagement using the anti-TIGIT antibody, the Mel1 sBC co-cultures displayed no significant differences compared to isotype antibody control cultures (**Figure 9K, dark grey and dark pink**), mimicking our results within the T cell co-culture experiments (**Figure 8F**). However, within the CD155 Mut sBC co-cultures, blockade of TIGIT signaling significantly increased NK cell-mediated sBC lysis (5:1: 2.67-fold, p=0.0038; 10:1: 2.47-fold, p<0.0001) compared to the isotype control group (**Figure 9K, light grey and light pink**). Collectively, these studies suggest that the expression of mutant CD155 on the surface of sBC has the ability to suppress both NK cell and T cell activation and sBC lysis, emphasizing the power of this system to inhibit both innate and adaptive immune responses.

## Discussion

The development of durable, pancreatic β-cell replacement therapies requires engineering strategies to enhance localized immune tolerance that can repress alloreactive and autoreactive T cell activation. The development of protocols to differentiate large pools of β-cells from hPSC have improved the ability to generate immunomodulatory cell populations (8, 11, 52, 53). Different approaches to accomplishing this goal have been attempted with mixed success. Preventing the expression of HLA molecules through deletion of *Beta-2 Microglobulin (β2M)* gene (HLA class I) or *class II transactivator (CIITA)* (HLA class II) genes proved attractive approaches to inhibit direct antigen presentation capabilities from the transplanted sBC (17–19, 54). However, a major limitation to this approach is HLA class I^-/-^ cells are targeted by NK cells through the “missing self” response (55, 56). Retention of HLA-E or HLA-G or expression of CD47 can limit the NK-mediated destruction of engineered cells (17, 57). We have previously engineered the expression of the check point inhibitor PD-L1 on sBC and demonstrated a reduction in stimulation of autoreactive T cell avatars (19). However, *β2M* knockout cell products may be fundamentally vulnerable in the event of tumorigenesis or exposure to infection due to challenges in immune surveillance. Therefore, approaches that can retain HLA expression while still effectively inhibiting T cell responses may be more attractive.

In this study, we demonstrated that sBC can be engineered to express CD155 to preferentially promote T cell co-inhibitory signaling through the CD155/TIGIT axis and restrain both NK cell and T cell activation. Notably, we demonstrate that compared to a WT CD155 variant, a higher-affinity mutant CD155 confers even greater protection from NK cell and antigen-specific T cell killing by reducing immune cell effector function. Our findings provide compelling evidence that CD155 modulation may serve as an effective immune evasion strategy to bolster long-term graft survival for β-cell replacement therapies in T1D. We have previously shown that the T1D-risk-associated CD226/TIGIT axis can be modulated to reduce autoreactive T cell responses by inhibiting co-stimulatory CD226 signaling; however, we demonstrate that this pathway can also be modulated by augmenting co-inhibitory TIGIT signaling to restrain autoimmunity and protect sBC. Furthermore, our data confirm that the membrane-bound CD155 Mut-expressing cells exhibit a higher binding efficiency for CD226 and an even greater binding for TIGIT compared to CD155 WT-expressing cells, as predicted by Matsuo et al (34).

Interestingly, Chimienti et al. have demonstrated that the genetic deletion of *CD155* in sBC in the context of HLA-I deficiency demonstrated the ability to circumvent the NK cell-targeted destruction (58). Hence, we hypothesize that the modulation of the CD155/TIGIT/CD226 axis in either direction – eliminating CD226:CD155 co-stimulatory signaling or bolstering TIGIT:CD155 co-inhibitory signaling – may enhance the protection of sBC from adaptive immune responses. To date, this is the first study to express CD155 on the surface of sBC, demonstrating its success as an approach to diminish NK cell and antigen-specific T cell activation through TIGIT-mediated co-inhibitory signaling. We also observed the ability for CD155-expressing sBC to suppress control antigen (MART-1) TCR avatars and polyclonal, allogeneic T cell proliferation, suggesting this mechanism may inhibit broad T cell engagement at the localized graft site following transplantation. Indeed, we corroborated that the enhanced protection previously conferred to CD155 Mut sBC was lost with the inclusion of an anti-TIGIT blocking antibody, emphasizing the role of CD155/TIGIT interactions in modulating immune responses (34). Notably, we did not observe the CD155 Mut sBC exhibiting greater susceptibility to CML from CD226:CD155 signaling during TIGIT blockade, relative to the Mel1 sBC, suggesting that CD226:CD155 signaling is not the driving signaling mechanism behind effector action in this system. Our group has recently demonstrated that inhibiting CD226 co-stimulatory signaling through anti-CD226 blocking antibodies inhibits murine effector T cell responses (28). Therefore, we would hypothesize that anti-CD226 treatment may yield additional protection to CD155-expressing sBC, however, additional studies of CD226:CD155 interactions in human sBC populations are needed.

Future studies should seek to assess the long-term and *in vivo* efficacy of engineering sBC to express immunomodulatory receptors, including whether CD155 engineering can be combined with other immunomodulatory approaches to enhance immune tolerance or Treg recruitment. Additionally, future pre-clinical studies should determine whether the protection conferred by CD155 expression differs as a result of TCR affinity, as our study employed 1E6 TCR avatars, which exhibit a lower TCR:pMHC affinity resemblant of the typical affinity of autoreactive T cells found *in vivo* (59). While impossible to assess accurately in existing pre-clinical models, more interrogation into the ability for CD155 expression to inhibit the polyclonal nature of a human *in vivo* immune response that includes both allogeneic and recurrent autoimmune responses is also necessary to determine the potential effectiveness in clinical practice. For this to become a reality, better model systems that can recapitulate the human autoimmune interaction with sBC are required (60, 61). Overall, our data support the continued investigation of engineering the expression of immunomodulatory molecules on sBC as a therapeutic strategy for enhancing the immune escape of sBC from innate immune and antigen-specific autoimmune T cell responses.

## Acknowledgements

We thank Andrew MacMillan-Ladd, Rachel Wilkes, and members of the Brusko and Russ Laboratories at the University of Florida Diabetes Institute for discussions and technical assistance. We also thank the UF Center for Immunology and Transplantation (CIT) for spectral flow cytometer and cell sorter access.

## Author Contributions

Conceptualization: MEB, JMB, TMB, HAR

Methodology: MEB, JMB, TMB, HAR

Investigation: MEB, JMB, MRP, JP

Formal Analysis: MEB, JMB

Visualization: MEB, JMB

Project Administration: MEB, JMB, TMB, HAR

Supervision: TMB, HAR

Funding Acquisition: TMB, HAR

Writing—original draft: MEB, JMB

Writing—r eview & editing: MEB, JMB, MRP, JP, TMB, HAR

## Data and Materials Availability

All data are available in the main text or the supplementary materials.

