## Supplemental Materials for "Human Stem Cell-Derived β-cells Expressing An Optimized CD155 Reduce Cytotoxic Immune Cell Function for Application in Type 1 Diabetes"

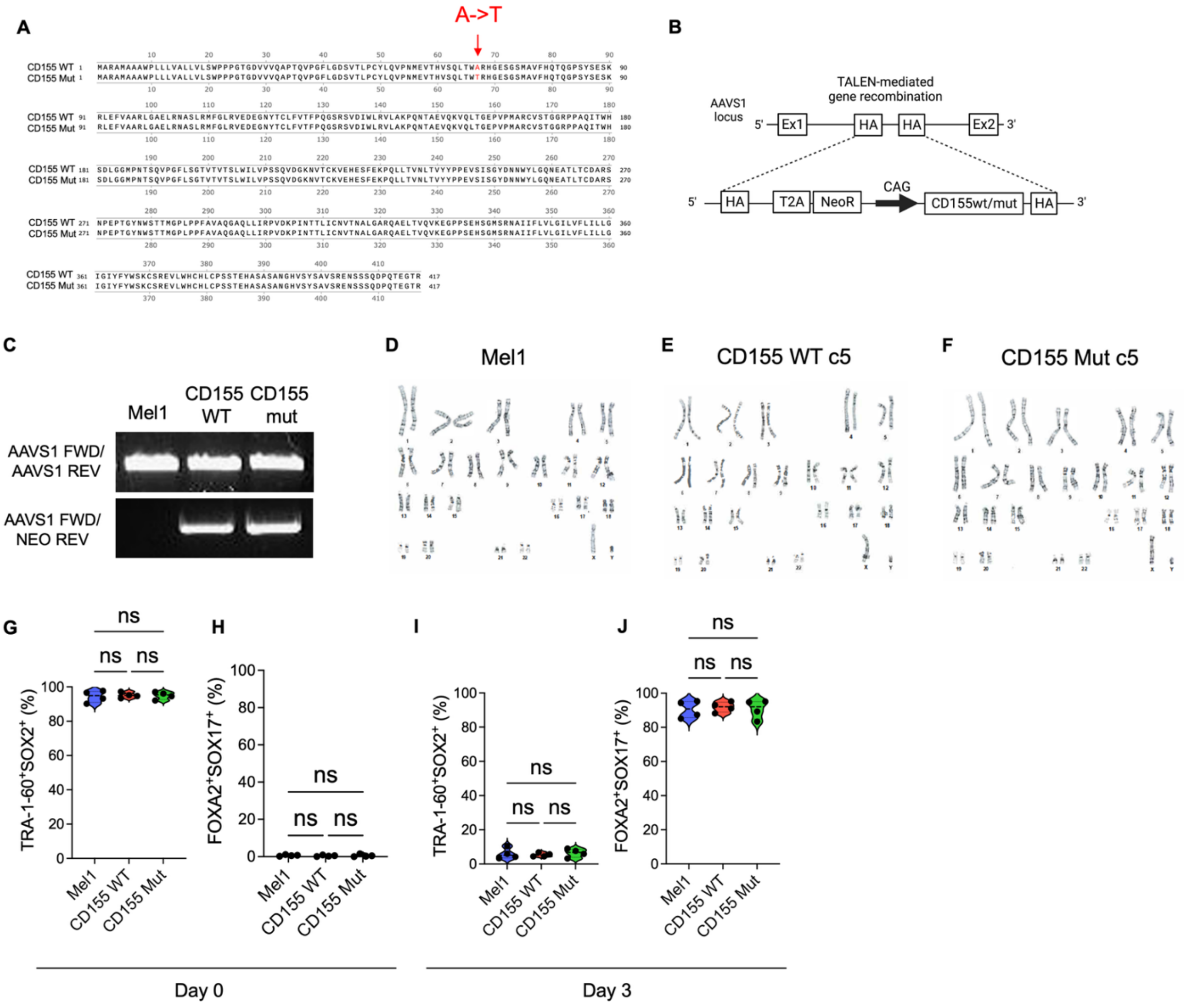


**Supplemental Fig. 1. Engineering and differentiation of CD155 edited hPSC.** (**A**) Amino acid sequence of WT CD155 compared to Mutant displaying the Ala67Thr mutation in red. (**B**) Schematic of gene overexpression construct of CD155 driven under the CAG promoter. (**C**) PCR Confirmation of gene insertion into the AAVS1 locus using amplification of the neomycin resistance gene. Representative image for G-banding metaphase karyotype analysis for (**D**) parental Mel1, (**E**) CD155 WT clone 5 (c5), and (**F**) CD155 Mut c5 hPSC demonstrating normal karyotyping in all three lines. (**G-J**) Flow cytometry of pluripotency markers TRA-1-60 and SOX2

(**G,I**) and definitive endoderm markers FOXA2 and SOX17 (**H,J**) in clusters on day 0 and day 3 of differentiation.

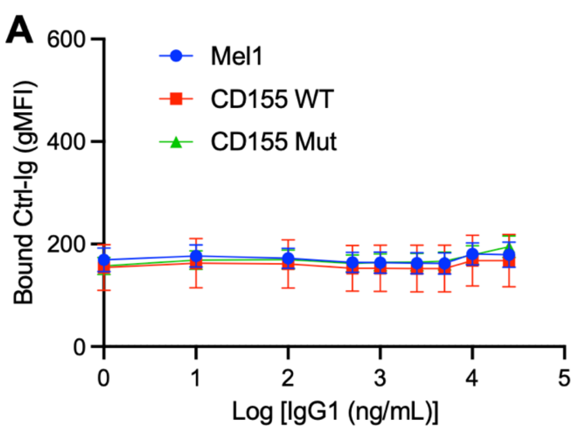


**Supplemental Fig. 2. CD155 expression does not influence binding to control proteins.** (**A**) hPSC or (**B**) sBC from the three cell lines were treated with fluorescently labeled IgG1 isotype control-Ig to compare binding efficiency between the parental Mel1 (blue), CD155 WT-expressing (red), and CD155 Mut-expressing (green) lines. Differences in Ig binding efficiency were characterized across a range of 0-25,000 ng/mL. Data reflects biological n=4/condition for hPSC and n=3/condition for sBC. No significant P-values were identified for one-way ANOVA with Bonferroni's multiple comparisons of AUC values between paired samples.

### Supplemental Tables:

#### Supplemental Table 1. Primers Used for Construct Design and Confirmation

| Application | Specification | Sequence (5' to 3') |
|---|---|---|
| Q5 Mutagenesis | Forward | GCTGACTTGGACGCGGCATGG |
| | Reverse | TGTGACACATGCGTCACCTC |
| CD155 Sequencing | Forward | AATTCGTCGACTGGATCCGG |
| | Reverse | CAGGAAACAGCTATGACCGC |
| | Internal | GGGTAGTACACGGTGAGGTT |
| Amplification with Homology Arms | Forward | TCGTATAGCATACATTATACGAAGTTATAATTCTGCAGATATCGGCCGGGAATTCGTCG |
| | Reverse | TGCATTCTAGTTGTGGTTTGTCCAAACTCATCAATGTATCTTAAGATCTGTTCAGGAAACAGCTATGACC |
| Line Confirmation | AAVS1 Forward | CCGGAACTCTGCCCTCTAAC |
| | AAVS1 Reverse | GTGGGCTTGTACTCGGCTAT |
| | Neomycin Reverse | GAACCTGCGTGCAATCCATC |

**Supplemental Table 2. Antibodies used for flow cytometry and immunofluorescence.**

| Target | Clone | Host | Fluorochrome | Concentration | Vendor | RRID |
|---|---|---|---|---|---|---|
| CD3 | OKT3 | Mouse | AF700 | 0.1 µg/mL | BioLegend | AB_2563408 |
| CD8 | SK1 | Mouse | AF700 | 0.2 µg/mL | BioLegend | AB_2562790 |
| CD27 | M-T271 | Mouse | Pacific Blue | 0.2 µg/mL | BioLegend | AB_2562505 |
| CD56 | QA17A16 | Mouse | Spark PLUS UV395 | 0.2 µg/mL | BioLegend | AB_3097564 |
| CD69 | FN50 | Mouse | BV711 | 0.2 µg/mL | BD Horizon | AB_2738443 |
| CD96 | NK92.39 | Mouse | BV421 | 0.2 µg/mL | BioLegend | AB_2629537 |
| CD112 | TX31 | Mouse | PE-Cy7 | 0.2 µg/mL | BioLegend | AB_2565732 |
| CD155 | SKII.4 | Mouse | AF647 | 0.2 µg/mL | BioLegend | AB_2721409 |
| CD226 | 11A8 | Mouse | PE-Cy7 | 0.2 µg/mL | BioLegend | AB_2616645 |
| CD274 | 29E.2A3 | Mouse | BV711 | 0.2 µg/mL | BioLegend | AB_2565764 |
| CD314 | 1D11 | Mouse | PE | 0.2 µg/mL | BioLegend | AB_492960 |
| HLA-A2 | BB7.2 | Mouse | Pacific Blue | 0.1 µg/mL | BioLegend | AB_2561569 |
| HLA-A,B,C | W6/32 | Mouse | PE | 0.1 µg/mL | BioLegend | AB_314875 |
| HLA-DR | L243 | Mouse | PerCP-Cy5.5 | 0.2 µg/mL | BioLegend | AB_893567 |
| HLA-DR | L243 | Mouse | BV570 | 0.2 µg/mL | BioLegend | AB_2650882 |
| Rat CD2 | OX-34 | Mouse | PE | 0.4 µg/mL | BioLegend | AB_2073811 |
| TIGIT | MBSA43 | Mouse | PerCP-eFluor710 | 0.2 µg/mL | Invitrogen | AB_10854428 |
| TRA-1-60 | TRA-1-60-R | Mouse | AF647 | 0.5 µg/mL | BioLegend | AB_1227812 |
| SOX2 | 14A6A34 | Mouse | AF594 | 5 µg/mL | BioLegend | AB_2562246 |

| | | | | | | |
|---|---|---|---|---|---|---|
| SOX17 | P7-969 | Mouse | AF488 | 0.5 µg/mL | BD Biosciences | AB_10893402 |
| FOXA2 | N17-280 | Mouse | PE | 0.25 µg/mL | BD Biosciences | AB_10716057 |
| C-peptide | C-PEP-01 | Mouse | AF488* | 0.83 µg/mL | Origene | AB_3712510 |
| CD155 | SKIL4 | Mouse | BV510 | 0.25 µg/mL | BioLegend | AB_2810528 |
| Insulin | Polyclonal | Guinea Pig | N/A | 5 µg/mL | DAKO | N/A<br>Cat # A0564 |
| NKX-6.1 | F55A10 | Mouse | N/A | 2 µg/mL | DSHB | AB_532378 |
| Mouse IgG | polyclonal | Donkey | AF647 | 0.5 µg/mL | ThermoFisher | N/A<br>Cat # A31571 |
| Guinea pig IgG | Polyclonal | Donkey | AF488 | 0.5 µg/mL | ThermoFisher | N/A<br>Cat # A11073 |

* Conjugated in-house

**Supplemental Table 3. LEGENDplex Supernatant Dilutions and Detection Range**

| LEGENDplex Assay Supernatant Dilutions | | |
|---|---|---|
| **Analyte** | **Dilution Factor** | **Adjusted Detection Range (pg/mL)** |
| IL-17A | 1; ND | 2.9 to 12,000 |
| IL-2 | 1; ND | 16.6 to 68,000 |
| IL-4 | 1; ND | 3.9 to 16,000 |
| IL-10 | 1; ND | 3.7 to 15,000 |
| IL-6 | 1; ND | 4.6 to 19,000 |
| TNF | 1; ND | 3.9 to 16,000 |
| Fas | 1; ND | 14.9 to 61,000 |
| FasL | 1 | 2.4 to 10,000 |
| IFN-γ | 100 | 390 to 1,600,000 |
| Granzyme A | 100 | 630 to 2,600,000 |
| Granzyme B | 1 | 14.1 to 58,000 |
| Perforin | 1 | 2.9 to 12,000 |
| Granulysin | 1 | 11.0 to 45,000 |
